# Cryo-EM Structure of Mycobacteriophage D29 Reveals a Specialized Portal-Capsid Junction and a Unique 13-Subunit Baseplate Collar

**DOI:** 10.64898/2026.11.10.765797

**Authors:** Biao Zhou, Zimu Li, Buhari Yusuf, Yi Zhao, Daming He, Zelong Huang, Luokai Wang, Mei Li, Xiangkai Zhen, Ji Gan, Tianyu Zhang, Liang Chen, Jinxing Hu, Xuemei Zeng, Xiaoli Xiong

**Affiliations:** State Key Laboratory of Respiratory Disease, Guangzhou Chest Hospital, Institute of Tuberculosis, Guangzhou Medical University, Guangzhou 510095, China; GMU-GIBH Joint School of Life Sciences, Guangdong Provincial Key Laboratory of Protein Modification and Disease, The Guangdong-Hong Kong-Macao Joint Laboratory for Cell Fate Regulation and Diseases, Guangzhou Medical University, Guangzhou, 511436, China; Guangzhou National Laboratory, Guangzhou 510005, China; State Key Laboratory of Respiratory Disease, CAS Key Laboratory of Regenerative Biology, Guangdong Provincial Key Laboratory of Stem Cell and Regenerative Medicine, Guangdong Provincial Key Laboratory of Biocomputing, Guangzhou Institutes of Biomedicine and Health, Chinese Academy of Sciences, Guangzhou 510530, China; University of Chinese Academy of Sciences, Beijing 100049, China; School of Clinical and Basic Medical Sciences, Shandong First Medical University & Shandong Academy of Medical Sciences, Jinan 250117, China; Biomedical Research Center of South China, College of Life Sciences, Fujian Normal University, Fuzhou 350117, China; Guangzhou Eighth People’s Hospital, Guangzhou Medical University, No.8 Huaying Road, Guangzhou, Guangdong, 510440, China; Key Laboratory of Microbial Pathogenesis and Interventions of Fujian Province University, the Key Laboratory of Innate Immune Biology of Fujian Province, College of Life Sciences, Fujian Normal University, Fuzhou, 350117, China

**Author notes:** Correspondence (T.Z.); (J.H.) (L.C.); (X.Z.); (X.X.). These authors contributed equally: Biao Zhou, Zimu Li, Buhari Yusuf, Yi Zhao.

## Abstract

Mycobacteriophages are promising agents against drug-resistant mycobacterial infections, yet their structural organization and host-recognition mechanisms remain poorly understood. Here, we determined cryo-EM structures of the capsid, connector, tail tube, and baseplate of the lytic, broad-host-range mycobacteriophage D29, revealing a near-complete virion architecture. At the capsid-portal interface, five copies of the previously uncharacterized protein gp19.1 bridge adjacent capsid subunits around the dodecameric portal, potentially reinforcing the symmetry-mismatched vertex. At the baseplate, 13 copies of gp6 assemble into a collar surrounding the sixfold-symmetric gp23-gp27 core. Remarkably, this 6:13 symmetry mismatch is accommodated through six distinct gp6-gp27 interfaces involving 12 gp6 subunits, leaving one subunit largely disengaged from the core. This asymmetric organization produces a tilted, outward-flaring collar that presents an extensive surface with the potential to engage the host cell. Whole-cell binding and plaque-competition assays further implicate the gp6 collar and the flexible trimeric central fiber gp31 in host recognition. Together, these findings reveal distinct structural adaptations at symmetry-mismatched interfaces and provide a structural framework for understanding mycobacteriophage assembly and host engagement.

## Introduction

Tuberculosis and infections caused by nontuberculous mycobacteria remain difficult to treat because of prolonged drug regimens, emerging antimicrobial resistance, and the capacity of mycobacteria to persist in diverse physiological states^1,2^. Lytic mycobacteriophages are therefore of considerable interest as antibacterial agents, diagnostic tools, and platforms for engineering host specificity^3–5^. Although phage structures and infection mechanisms have been extensively characterized in several bacterial systems^6–9^, the molecular basis of mycobacteriophage assembly and host recognition remains comparatively underexplored^10,11^.

Most mycobacteriophages are tailed double-stranded DNA viruses within the class *Caudoviricetes* and display a siphovirus-like morphology with an icosahedral capsid, a portal-containing connector, a long non-contractile tail, and a distal baseplate^12–16^. Although these modules share a conserved architectural framework, recent cryo-electron microscopy (cryo-EM) studies have shown that the surface-exposed and distal components of mycobacteriophages are highly diversified. In Bxb1, an elaborate tail-tip cage and peripheral C-terminal-extension-like (CTE-like) domains participate in host-associated conformational remodeling^13^. In Douge, multiple Ig-like glycan-binding domains are distributed across the capsid, tail tube, and baseplate, generating an extensive host-interaction surface^16^. These structures establish that related long-tailed mycobacteriophages can employ fundamentally different strategies to contact the mycobacterial cell envelope.

Mycobacteriophage D29, first isolated in 1954^17^, is a lytic member of subcluster A2 closely related to the temperate mycobacteriophage L5^18^. Comparative genomic analyses suggest that D29 evolved from an L5-like temperate ancestor through an approximately 3.6-kb deletion encompassing the lysogeny repressor locus^19^, consistent with its strictly lytic lifestyle^18^. D29 infects both fast-and slow-growing mycobacteria^17,18^, including *Mycobacterium smegmatis* (Msm) and selected *Mycobacterium tuberculosis* (Mtb) strains^20–22^. Its broad host range and lytic activity have made D29 a valuable tool for mycobacterial genetics, reporter-phage diagnostics, and phage-based antimicrobial studies^23–26^. Nevertheless, despite its extensive genetic characterization and diverse applications, the structural basis of host recognition by D29 remains poorly understood.

Unlike Bxb1 and Douge^13,16^, D29 lacks the C-terminal extension (CTE) domains found in the Bxb1 capsid and tail tube^13^, as well as the widely distributed glycan-binding domains observed in Douge^16^. How D29 interacts with its host in the absence of these reported host-recognition structures remains unknown. Previous studies demonstrated that the minor tail protein gp6 of the related mycobacteriophage L5 binds mycobacterial cells and identified a homolog encoded by D29^27,28^. Fusion of L5 gp6 with antimicrobial peptides was also reported to enhance the elimination of intracellular Mtb in macrophages^29^. However, whether the D29 minor tail protein gp6 performs a comparable host-binding function, where it is located within the virion, and how it coordinates with other baseplate components remain unclear, as does the overall virion architecture of D29.

In this study, we determined cryo-EM structures of the major architectural modules of mycobacteriophage D29, including the capsid, connector, tail tube, and baseplate, and constructed a composite structural model of the whole virion. These structures identify gp19.1 as a previously uncharacterized component of the symmetry-mismatched portal-capsid junction and reveal an unusual 13-subunit gp6 collar surrounding the sixfold-symmetric baseplate core. Unlike the elaborate tail-tip assembly of Bxb1^13^ and the widespread Ig-like host-interaction modules of Douge^16^, D29 exhibits a distinct organization of its distal adsorption apparatus. Combined structural and functional analyses further implicate gp6 and the central fiber gp31 in host binding. Together, these findings reveal distinct structural adaptations at symmetry-mismatched interfaces and provide a framework for understanding mycobacteriophage assembly, host recognition, and future structure-guided engineering for therapeutic applications.

## Results

### Overall architecture of mycobacteriophage D29

D29 particles purified by sucrose density-gradient ultracentrifugation were analyzed by single-particle cryo-EM. Cryo-EM micrographs and reference-free two-dimensional class averages revealed a typical siphovirus-like morphology, comprising an icosahedral capsid, a neck complex, a long flexible non-contractile tail, and a distal baseplate (**Fig. 1a-b**). High-resolution reconstructions of the individual structural modules were assembled into a composite model of the mature D29 virion (**Fig. 1c-e and S1)**. The visible portion of the virion is ∼2240 Å in length, comprising an ∼1900 Å capsid-tail assembly and an ∼340 Å distal assembly formed by the baseplate and central fiber. The major capsid protein (MCP) gp17-formed icosahedral capsid is approximately 520 Å in diameter and encloses the densely packaged double-stranded DNA genome (**Fig. 1c-e and S1**).

**Fig. 1.**
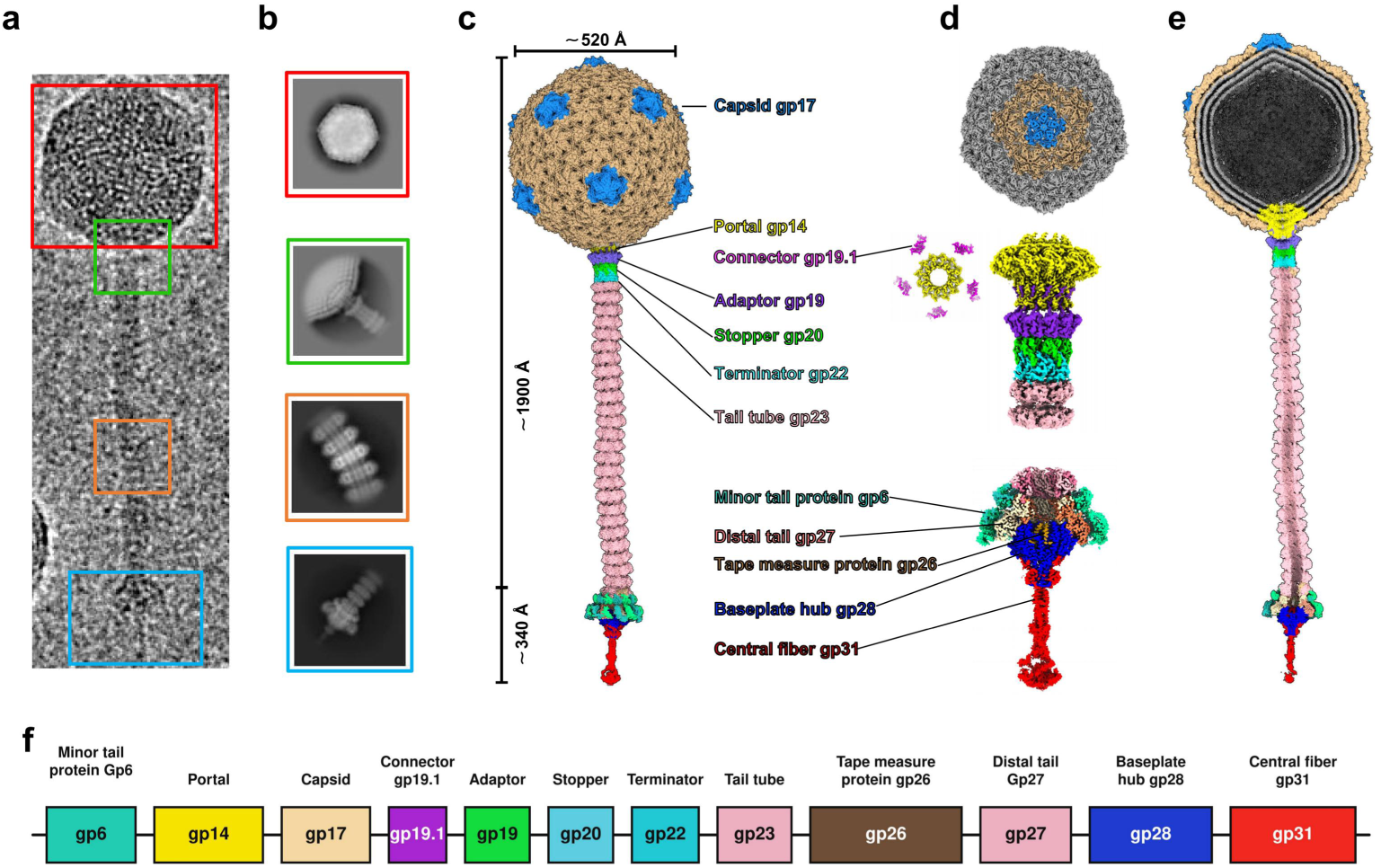
Overall architecture of mycobacteriophage D29. **a**, Cryo-EM micrograph of a D29 particle. **b**, 2D class averages of the D29 capsid, portal complex, tail tube and baseplate components. **c**, A composite cryo-EM map of the entire D29 particle, with density segments for each structural component labeled and colored as in (a). **d**, Exploded views of the portal-capsid junction, connector and baseplate, showing their component organization. **e**, Cross-sectional view of the composite cryo-EM map of the entire D29 particle. **f**, Genomic organization of the resolved D29 structural proteins.

At the unique 5-fold capsid vertex, the dodecameric portal protein gp14 connects axially to the dodecameric adaptor gp19, followed by the hexameric stopper gp20 and terminator gp22, together forming a continuous channel between the capsid and tail (**Fig. 1c-d**). By contrast, gp19.1 is positioned peripherally around the portal and laterally bridges adjacent gp17 capsid subunits at the portal-capsid interface (**Fig. 1d**). The ∼1000 Å long tail tube comprises approximately 25 stacked hexameric rings formed by the tail tube proteins (TTP), gp23 (**Fig. 1c-e**). The distal baseplate comprises multiple copies of the minor tail protein (MTP) gp6, distal tail protein (DTP) gp27, and baseplate hub protein gp28, together with densities assigned to residues 818-836 of the C-terminal domain of the tape measure protein (TMP) gp26 and the trimeric central fiber protein (CFP) gp31. Together, these components form a multicomponent apparatus involved in host recognition and attachment (**Fig. 1c-d**). We further mapped these structural assignments onto the D29 genome, linking each resolved virion component to its corresponding gene (**Fig. 1f**).

### The phage D29 Capsid

We reconstructed the D29 capsid to an overall resolution of 2.5 Å by imposing icosahedral symmetry (**Fig. S1a and S2**). The capsid adopts a T = 7 icosahedral lattice and is primarily assembled from the MCP gp17. gp17 forms 60 hexameric capsomers (hexons) and 11 pentameric capsomers (pentons), while the twelfth 5-fold vertex is occupied by a specialized multicomponent portal complex. The pentons are located at the vertices of the icosahedron and are surrounded by five neighboring hexons (**Fig. 1a**). The maximum spans of the pentons and hexons are approximately 143 Å and 168 Å, respectively (**Fig. S3a-b**). Accordingly, each icosahedral asymmetric unit contains 11 quasi-equivalent gp17 subunits, six of which form a complete hexon, whereas the remaining five form a complete penton (**Fig. 2b**).

**Fig. 2.**
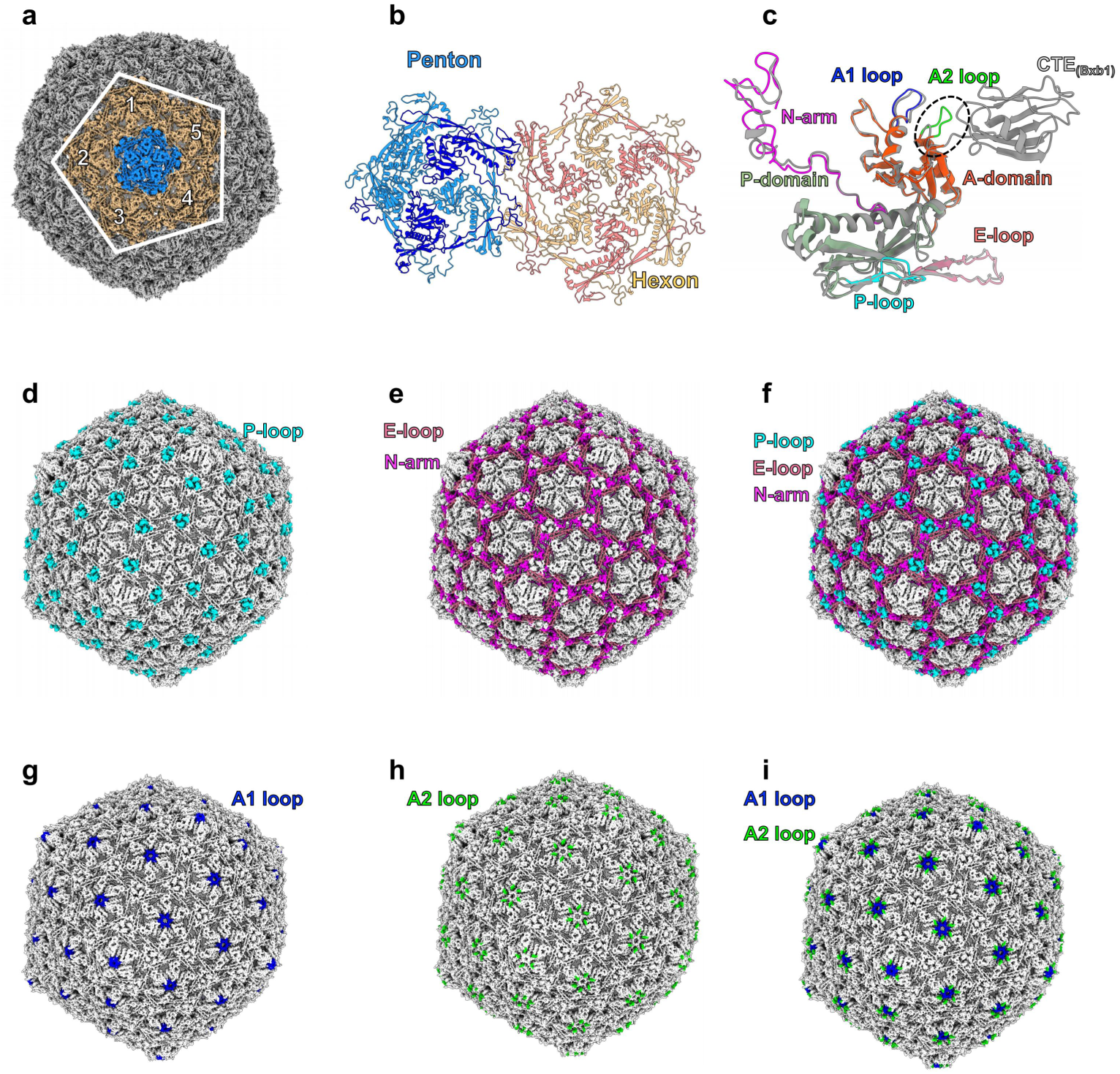
Structure and organization of the D29 capsid. a, Cryo-EM reconstruction of the capsid, highlighting five hexons surrounding a central penton. **b**, Cartoon representation of a representative penton-hexon pair, comprising five and six gp17 subunits, respectively. **c**, Structural superposition of D29 gp17 and Bxb1 major capsid protein, highlighting the conserved N-arm, P domain, A domain, E-loop, and P-loop, as well as the A1 loop and D29-specific A2 loop. The Bxb1 major capsid protein is shown in gray. **d-f**, Capsid surface views highlighting the P-loops (**d**), E-loops and N-arms (**e**), and their combined distribution (**f**), which forms an interwoven network across the capsid lattice. **g-i**, Capsid surface views showing the A1 loops (**g**), A2 loops (**h**) and their combined concentric arrangement around the capsomer axes (**i**).

The D29 MCP gp17 adopts the canonical HK97 fold, a highly conserved capsid-protein architecture widely found in tailed double-stranded DNA bacteriophages and herpesviruses^30–32^. Each gp17 subunit consists of an N-terminal arm (N-arm; residues 1-49), an extended E-loop (residues 55-90), a peripheral P domain (residues 91-149 and 246-304), and an axial A domain (residues 150-245 and 305-311) (**Fig. 2c**). Although gp17 subunits in the pentons and hexons share the same conserved structural core, their N-arms, E-loops, and P-loops adopt distinct conformations, enabling the MCP to accommodate the different local curvatures and quasi-equivalent environments of the pentons and hexons (**Fig. S3c**).

Next, we performed a DALI search, which identified Bxb1 MCP as its closest structural homolog, with a root-mean-square deviation (RMSD) of 1.00 Å (**Fig. 2c**). Although D29 MCP gp17 lacks the Ig-like CTE domain of Bxb1 MCP^13^, the two proteins share a conserved HK97-fold core, with closely superimposable A and P domains and E-loops (**Fig. 2c**). By contrast, the conserved P-loops form trimeric tower-like structures at the 3-fold axes (**Fig. 2d**), where they interact with three neighboring E-loops and an N-arm trimer to generate a chainmail-like capsid network (**Fig. 2e-f and S3d**). These contacts include a hydrogen bond between P-loop N283 and E-loop K78, together with extensive hydrophobic interactions among two N-arms and one E-loop (**Fig. S3e-f**), thereby stabilizing the D29 capsid.

Compared with the Bxb1 MCP, a distinctive feature of D29 MCP gp17 is an additional A2 loop (_155_ADTTGATT_162_) in the A domain (**Fig. 2c**). At the centers of pentons and hexons, the A2 loops forms an outer ring surrounding the A1 loops (_199_KDQNGRP_205_, **Fig. 2g-i and S3g-i**). Both loops contain polar, solvent-exposed residues and create concentric hydrophilic structures around the capsomer opening (**Fig. 2g-i and S3g-j**), potentially reinforcing the capsomer axis and maintaining capsid integrity.

Cryo-EM micrographs revealed a subset of particles with markedly reduced internal density, and the corresponding 2D class averages showed intact capsid shells without discernible packaged DNA (**Fig. S4a**). We subsequently reconstructed a structure of the genome-free capsid at 4.2 Å overall resolution (**Fig. S1d and S2g**). The structure retained an intact icosahedral shell surrounding a largely empty interior (**Fig. S4b**). Compared with the genome-packed capsid, the genome-free capsid exhibited a larger diameter (**Fig. S4c-d**), suggesting that genome release is accompanied by expansion or relaxation of the capsid shell. Similar post-ejection structural changes have also been reported for Bxb1 and Douge^13,16^.

### Gp19.1 reinforces the symmetry-mismatched portal-capsid junction

To resolve the portal-capsid junction, particles from the icosahedral capsid reconstruction were subjected to symmetry expansion, duplicate removal, volume alignment, and 3D classification. Following non-uniform refinement without imposed symmetry (C1), the capsid density was clearly resolved, whereas the portal structure was poorly resolved (**Fig. S1a and S5a-c**). This observation suggests heterogeneity in the spatial relationship between the capsid and the portal complex. The capsid-portal particles were further refined with C5 and C6 symmetry, yielding reconstructions at overall resolutions of 2.9 and 2.8 Å, respectively (**Fig. S1a and S5d-i**). The C5 map resolved the five capsid hexons surrounding the portal, including additional densities between adjacent hexons that were subsequently assigned to gp19.1, whereas the portal density remained poorly resolved (**Fig. S5d-f**). Conversely, the C6 map resolved the connector complex, whereas the surrounding capsid density was poorly resolved (**Fig. S5g-i**). The two maps were combined for structural visualization (**Fig. 3a-b and S1a**).

**Fig. 3.**
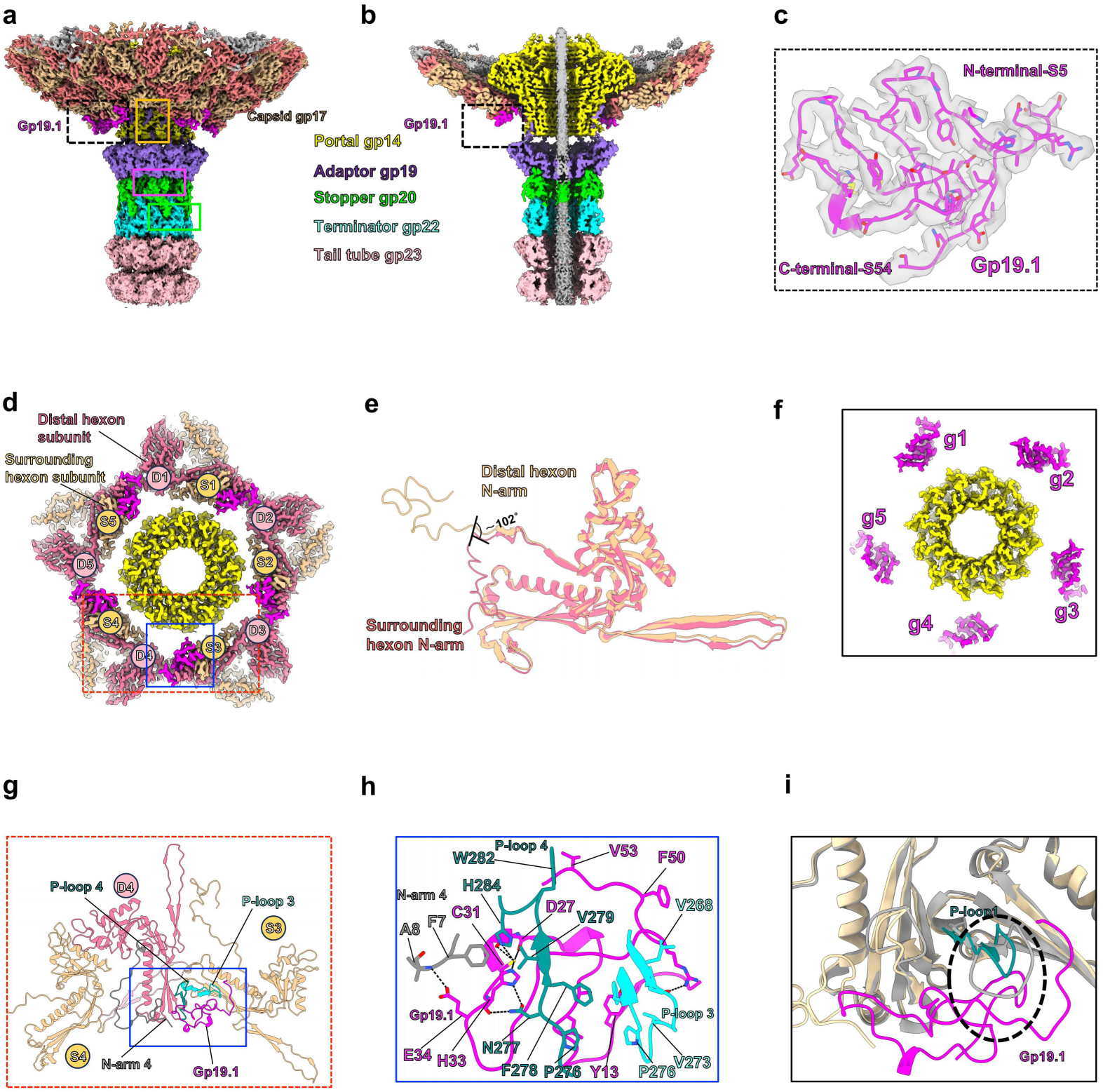
Architecture of the D29 portal-capsid junction. **a**, Side view of the portal vertex and connector complex, with the constituent proteins shown in different colors. **b**, Cutaway view showing the continuous channel through the gp14 portal, gp19 adaptor, gp20 stopper, gp22 terminator and gp23 tail tube. **c**, Fit of the gp19.1 atomic model (residues 5-54) into the cryo-EM density. **d**, View from inside the capsid showing the C12 gp14 portal, five surrounding gp17 hexons and five gp19.1 molecules. Subunits from the surrounding and distal hexons are designated S1-S5 and D1-D5, respectively. **e**, Superposition of representative gp17 subunits from the surrounding and distal hexons, showing the different orientations of their N-arms. **f**, Cryo-EM map showing five gp19.1 molecules surrounding the gp14 portal. **g**, Cartoon representation of a gp19.1 molecule positioned between neighboring gp17 subunits. **h**, Cartoon representation of the interactions between gp19.1 and the P-loops and N-arm of adjacent gp17 subunits. Hydrogen bonds are indicated by dashed lines. **i**, Structural superposition of gp17 subunits surrounding the portal vertex and a conventional fivefold vertex (gray), revealing a steric clash between the P-loop at the conventional vertex and gp19.1 (black dashed circle).

As in other tailed phages^33,34^, the pentameric capsomer at the unique bottom fivefold vertex of the D29 capsid is replaced by a dodecameric gp14 portal. Together, the portal protein gp14, adaptor gp19, stopper gp20, and terminator gp22 form the connector complex, which links the capsid to the tail tube and forms a channel for controlled genome ejection (**Fig. 3a-b**). The incorporation of the 12-mer gp14 portal into the C5 capsid vertex creates an intrinsic symmetry mismatch at the portal-capsid junction. The dodecameric gp14 portal is surrounded by five gp17 hexons, each contributing two subunits to the portal-proximal region. This arrangement creates a symmetry mismatch between the C12 portal and the C5 organization of the surrounding capsid, involving 12 gp14 protomers and 10 gp17 subunits (**Fig. 3d**). The ten gp17 subunits can be divided into two groups according to their positions relative to the portal: S1-S5 (S for surrounding), which form an inner ring adjacent to the gp14 protomers, and D1-D5 (D for distal), which are positioned further from the portal (**Fig. 3d**). Notably, the N-arms of the S and D subunits adopt markedly different orientations, with a relative angular difference of approximately 102° (**Fig. 3e**).

Interestingly, five additional protein densities are observed at the interfaces between adjacent gp17 hexons surrounding the portal vertex (**Fig. 3b-d**). Based on their local structural features and atomic-model fitting, these densities were assigned to the small phage protein gp19.1. Gp19.1 comprises 64 amino acids, of which residues 5-54 could be reliably modeled, whereas the remaining N- and C-terminal residues were unresolved (**Fig. 3c**). The five gp19.1 molecules are arranged in a discontinuous 5-fold-symmetric ring surrounding the gp14 portal complex (**Fig. 3f**). Thus, gp19.1 follows the fivefold organization of the surrounding capsid rather than the twelvefold symmetry of the portal.

Each gp19.1 molecule is located between two adjacent gp17 hexons and bridges two S subunits and one D subunit (**Fig. 3d and 3g**). Specifically, gp19.1 interacts with the P-loops of one S subunit and the D subunit, as well as the N-arm of the other S subunit, through hydrogen bonds and hydrophobic contacts (**Fig. 3h**). These interactions establish a local inter-hexon bridge that links three gp17 subunits at the symmetry-mismatched vertex. Structural comparison with the capsid subunits surrounding the Bxb1^13^ portal revealed a more open arrangement of gp17 subunits around the D29 portal, characterized by greater separation between neighboring subunits and the formation of local interfacial gaps (**Fig. S6a**). The five gp19.1 molecules occupy these gaps and connect adjacent hexons, potentially compensating for their relatively loose packing (**Fig. S6b**). This organization suggests that gp19.1 acts as a molecular brace that reinforces the capsid framework surrounding the symmetry-mismatched portal.

Unlike capsid decoration proteins distributed across repeated capsid sites or around 3-fold axes in other phages^35–37^, gp19.1 binding is confined to the unique portal-capsid junction in D29. Structural superposition of the capsid regions surrounding conventional fivefold vertices and the portal vertex revealed that gp19.1, when placed at the equivalent position, would sterically clash with the N-arms and P-loops of D29 MCPs at conventional vertices (**Fig. 3i**). These observations suggest that gp19.1 selectively recognizes local capsid structural features associated with the capsid-portal junction. Phylogenetic analysis further identified gp19.1 homologs across multiple mycobacteriophage clades, suggesting that related proteins may serve similar structural roles (**Fig. S6c**).

Together, these findings reveal a distinctive structural adaptation to the capsid-portal symmetry mismatch in D29. Five gp19.1 molecules bridge the interfaces between neighboring MCP hexons surrounding the dodecameric portal, potentially providing additional stabilization at the capsid-portal vertex. Although the resolved regions of gp19.1 do not contact the portal, contacts involving its unresolved termini cannot be excluded. Unlike the capsid-portal interfaces of mycobacteriophages Bxb1^13^ and Douge^16^, that of D29 could not be fully resolved. This may reflect relative motion between the capsid and the portal complex. The recruitment of the accessory protein gp19.1 may represent a strategy for stabilizing this dynamic, symmetry-mismatched interface.

### Connector

Similar to the connectors of other long-tailed phages^13,16^, the D29 connector comprises a dodecameric portal gp14-adaptor gp19 subcomplex, followed by hexameric rings of stopper gp20 and tail terminator gp22. The assembly spans approximately 210 Å and mediates the transition from the capsid to the phage tail tube, which is composed of stacked hexameric rings (**Fig. 4a**). Portal gp14 adopts the conserved mushroom-shaped architecture, with an outer diameter of approximately 165 Å and a minimum channel diameter of 32 Å (**Fig. 4b**). It has a negatively charged lumen that has been proposed to facilitate dsDNA translocation^38,39^.

**Fig. 4.**
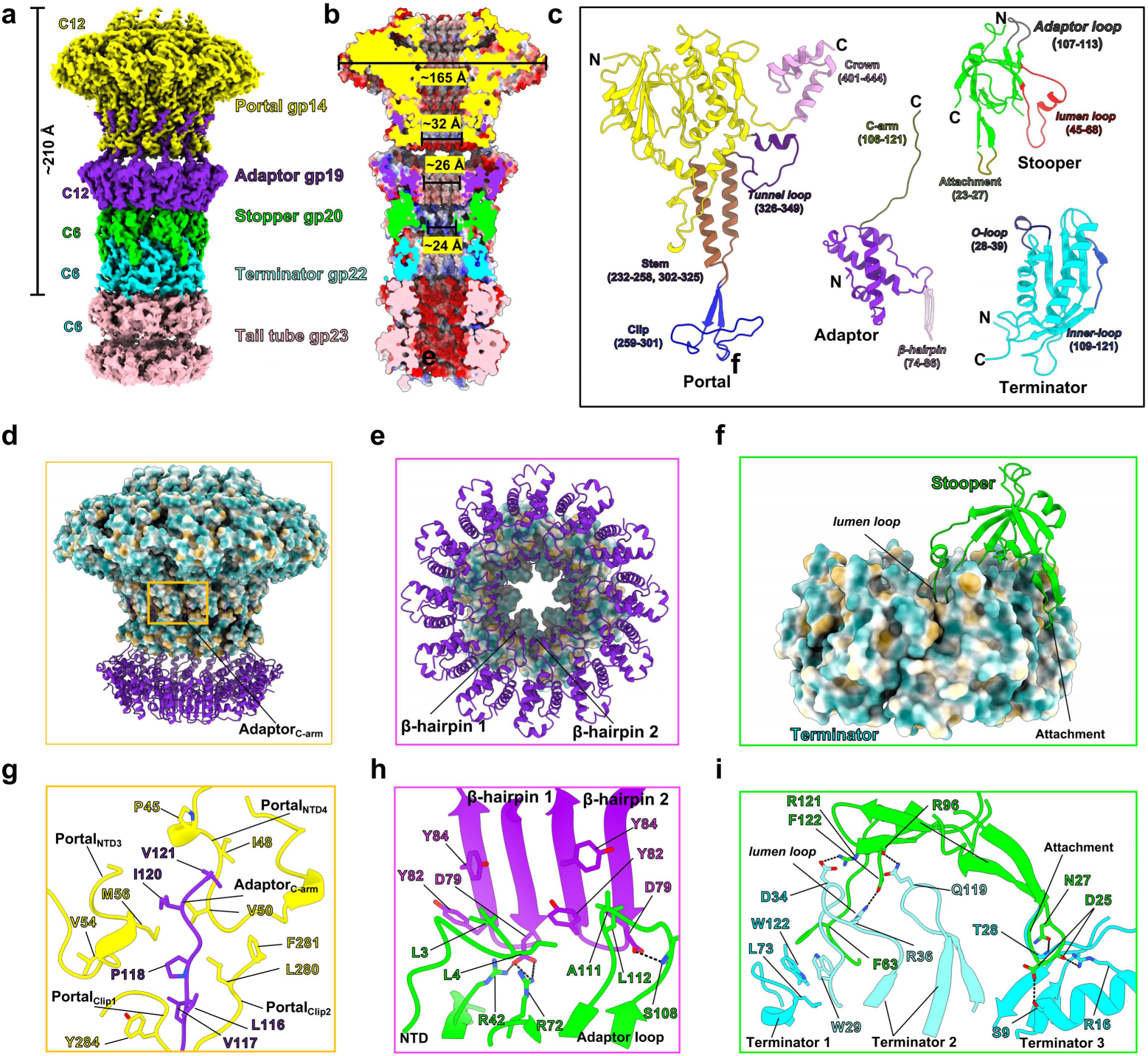
Structure and inter-subunit interactions of the D29 connector. **a**, Cryo-EM map of the D29 connector, comprising the C12 gp14 portal and gp19 adaptor, followed by the C6 gp20 stopper, gp22 terminator and gp23 tail tube. **b**, Cutaway view of the connector colored by electrostatic surface potential, showing the central channel and its indicated diameters. **c**, Cartoon representations of gp14, gp19, gp20 and gp22, with their structural elements indicated. **d**, Interaction between the gp19 adaptor C-arm and the gp14 portal. **e**, Top view of the gp19 adaptor dodecamer, showing the two β-hairpins from each subunit forming a 24-stranded β-barrel. **f**, Interaction between the gp20 stopper and gp22 terminator. **g-i**, Cartoon representations of the interactions at the gp14-gp19 (g), gp19-gp20 (h) and gp20-gp22 (i) interfaces. Hydrogen bonds are indicated by dashed lines.

The C-terminal arm of each adaptor gp19 inserts between the clip domains of two adjacent portal subunits, extending their β-sheets and forming additional hydrophobic contacts (**Fig. 4d and 4g**). The adaptor gp19 subunits assemble into a dodecameric ring containing a central β-barrel, which continues the portal channel (**Fig. 4e**). Beneath the adaptor, β-hairpins from stopper gp20 contact the N-terminal regions and adjacent loops of gp19 through hydrogen-bonding and hydrophobic interactions (**Fig. 4h**). Each stopper gp20 subunit subsequently contacts three neighboring gp22 subunits through its lumen-facing and attachment loops (**Fig. 4f and 4i**). These distributed interactions anchor the hexameric terminator ring to the stopper and connect the connector to the first tail-tube ring. Together, these components form a continuous channel and mediate the structural transition from the 12-fold symmetric portal-adaptor assembly to the 6-fold symmetric stopper-terminator and tail-tube complex. This overall organization resembles the connectors of phages NF5^40^, lambda^41^, SPP1^42^, Chi^43^, ΦXacJX1^44^ and other long-tailed phages, although the D29 portal-adaptor interface is primarily concentrated around the extended gp19 arms. In genome-free D29 particles, the structure of the “portal-adaptor-stopper-terminator” connector assembly remains largely unchanged, whereas densities corresponding to dsDNA and the TMP are absent (**Fig. S7**). Although these particles were not generated by experimentally induced genome ejection, the absence of these axial densities is consistent with TMP release or disordering, leaving an open connector-tail channel without substantial structural rearrangement of the connector assembly^45,46^.

### Tail tube

Mycobacteriophage D29 possesses a ∼1000 Å-long tail, composed of 25 layers of stacked TTP hexameric rings that form a tube with a diameter of 104 Å and a stacking spacing of 35 Å (**Fig. S8b-d**). The tail tube structure was resolved at 3.2 Å resolution by averaging particles spanning four tail tube ring layers, with refinement focused on the two central layers (**Fig. S2b**). Within the tail tube region (TTR), densities corresponding to TMP are observable, with an approximate diameter of 17 Å (**Fig. S8c-d**). In D29, each TTP subunit comprises a β-core formed by eight β-strands, an outer loop (residues 23-50), a protruding β-hairpin (residues 68-86) contributed by the β2-β3 region, and a buried α-β loop (residues 101-128) (**Fig. S8a**). Unlike Bxb1 and Douge^13,16^, D29 TTP lacks an exposed CTE domain (**Fig. S8a**).

At each inter-ring interface, the hinge loop projects downward into a hydrophobic groove formed by the β-sheet region of a TTP subunit in the adjacent lower ring, with the interaction supported primarily by hydrogen bonds and hydrophobic contacts (**Fig. S8e-f**). Superposition of the six subunits within one ring layer reveals a highly static core fold (RMSD, 0.442 Å), whereas their hinge loops adopt distinct conformations and orientations (**Fig. S8g**). Similar conformational variations in hinge loops mediating inter-ring interactions have been observed in Bxb1, where they accommodate relative tilting between adjacent ring layers^13^. By analogy, the flexible hinge loops of D29 TTP, which connect successive tail-tube rings, likely accommodate local geometric variations between adjacent rings, thereby preserving structural continuity while allowing mechanical flexibility.

### Baseplate

Phage tail structures are critical determinants of host specificity^47^. The baseplate, located at the distal end of the tail tube, mediates host recognition during phage infection^13,48,49^. Reconstruction of the complete baseplate with C1 symmetry yielded a 3.3 Å resolution map, revealing the peripheral regions, albeit at limited local resolutions (**Fig. S1c**). Within this reconstruction, the tail-tip density was only partially resolved but suggested a trimeric organization (**Fig. S1c**). Guided by this observation, we imposed C3 symmetry to improve the resolution of the tail tip, yielding a 3.16 Å reconstruction with enhanced density continuity that enabled the trimeric region of the tail-tip central fiber to be resolved (**Fig. S1c and 5a**).

Integration of these complementary reconstructions revealed that the D29 baseplate is approximately 210 Å in diameter and that its visible portion is approximately 340 Å long, with an overall architecture reminiscent of a desk lamp (**Fig. 5a-b**). Structural modeling of the C1 map defined the spatial arrangement of six constituent proteins within this assembly (**Fig. 5e**). Its central core comprises a hexameric TTP gp23 ring (**Fig. 5f and 5j**), a two-tiered DTP gp27 hexamer directly beneath it (**Fig. 5g and 5k**), and a trimeric gp28 hub at the distal end (**Fig. 5i and 5m**). The upper tier of gp27 contacts gp23 (**Fig. 5g and 5k**), whereas its lower tier expands outward. Within the central channel, the C-terminal regions of three TMP gp26 subunits lie inside the gp28 hub (**Fig. 5h and 5l**), which anchors an elongated trimeric CFP gp31 fiber projecting distally (**Fig. 5a-b**). Surrounding the gp23-gp27 core is a tilted, lampshade-like gp6 collar that forms the outermost layer of the baseplate (**Fig. 5c-d**). The collar comprises 13 subunits, as resolved in the C1 reconstruction and supported by representative 2D class averages (**Fig. S9**). Its upper region forms a well-ordered ring with approximate 13-fold rotational symmetry (**Fig. S10a-c**), whereas its lower region lacks clear rotational symmetry, with individual subunits adopting different orientations and exhibiting substantially lower local resolution (**Fig. S10d**). The exposed, flexible lower region of the gp6 collar and the outward-projecting regions of gp27 may both contribute to host recognition and attachment, consistent with the host-binding activities of these proteins described below.

**Fig. 5.**
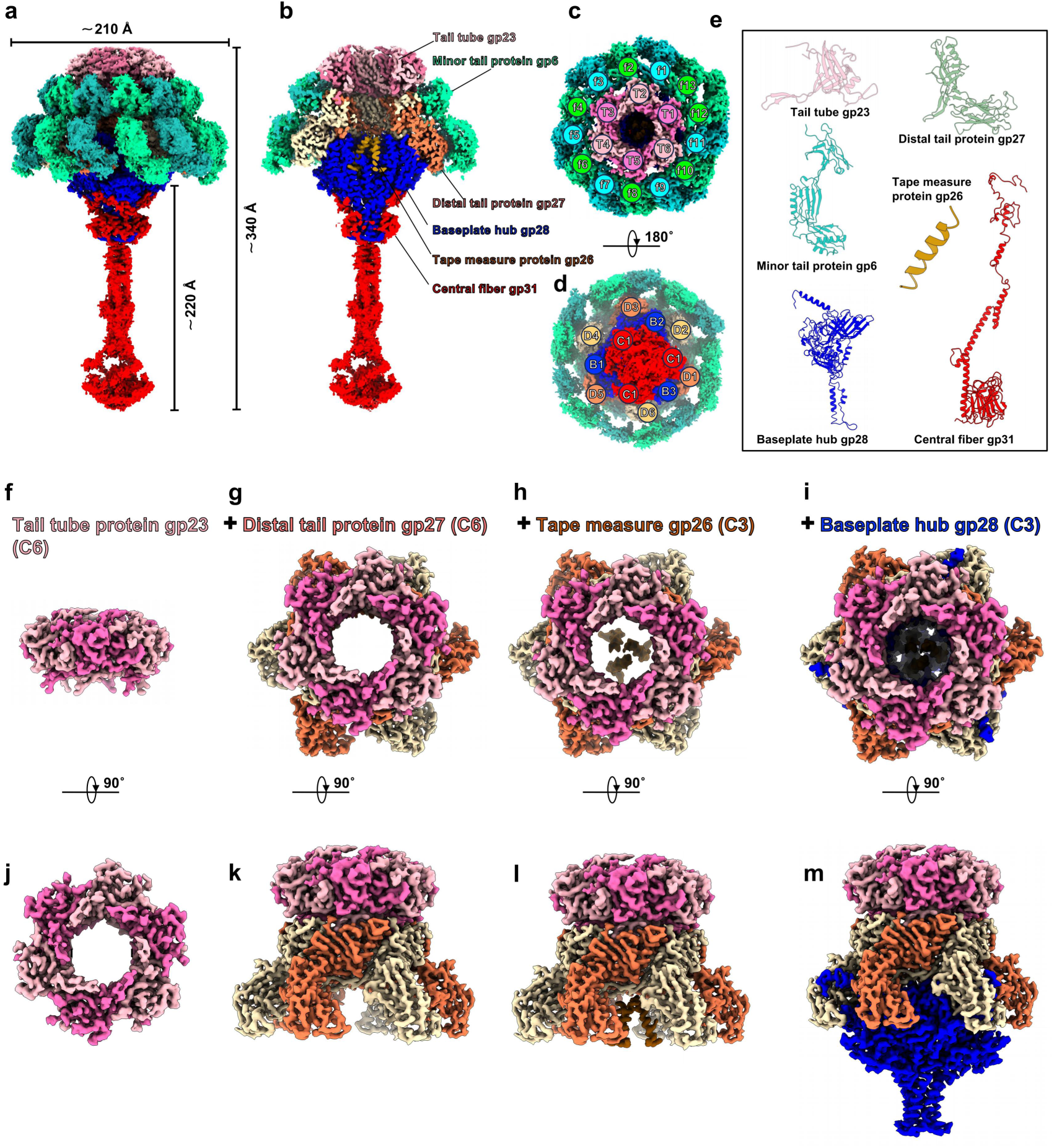
Overall architecture of the D29 baseplate. **a**, Cryo-EM map of the complete D29 baseplate, with the indicated dimensions. **b**, Cutaway view showing the spatial organization of gp23, gp6, gp27, gp26, gp28 and gp31. **c**, View from the tail-tube side, showing the C6 gp23 ring (T1-T6) surrounded by the 13-copy gp6 collar (f1-f13). **d**, View from the distal side, showing the organization of the C6 gp27 ring (D1-D6), C3 gp28 hub (B1-B3) and trimeric gp31 central fiber (C1-C3). **e**, Cartoon representations of the individual baseplate proteins. **f-i**, Axial views showing the progressive addition of the C6 gp27 ring (**g**), C3 gp26 (**h**) and C3 gp28 hub (**i**) to the C6 gp23 tail-tube ring (**f**). **j-m**, Corresponding side views of the structures shown in **f-i**, respectively.

Although the overall desk-lamp-like morphology of the D29 baseplate resembles the distal adsorption complexes of other long-tailed phages, including mycobacteriophages Douge^16^ and Bxb1^13^, lactococcal siphophages p2^50^ and TP901-1^51^, and Brochothrix phage NF5^40^, their underlying symmetry organizations differ substantially. These phages generally exhibit transitions between 6-fold-symmetric central assemblies and 3-fold-symmetric distal components, with peripheral structures displaying distinct oligomeric arrangements. In contrast, the D29 baseplate exhibits an unusual 6:13 symmetry mismatch between the conserved C6-symmetric gp23-gp27 central core and the surrounding 13-subunit gp6 collar, representing a distinctive architectural organization among mycobacteriophages.

### Symmetry-mismatched contacts between the 13-copy gp6 collar and the C6 gp23-gp27 core

Given the unusual baseplate architecture, we examined how the 6:13 symmetry mismatch between the C6-symmetric gp23-gp27 central core and the surrounding 13-copy gp6 collar is accommodated. The gp23 hexamer is positioned within the upper region of the gp6 collar (**Fig. 6a**), with only its lower rim extending into the central opening of the upper pseudo-C13 gp6 ring (**Fig. 6e**). The gp6 collar is tilted relative to the gp23-gp27 core, resulting in uneven spacing between individual gp23 subunits and the surrounding gp6 ring. Consequently, gp23 subunits T1-T3 (**Fig. 6b-d**) lie closer to the gp6 ring than T4-T6 (**Fig. 6f-h**). Despite this spatial proximity, no specific interactions are observed between the gp23 hexamer and the upper pseudo-C13 gp6 ring.

**Fig. 6.**
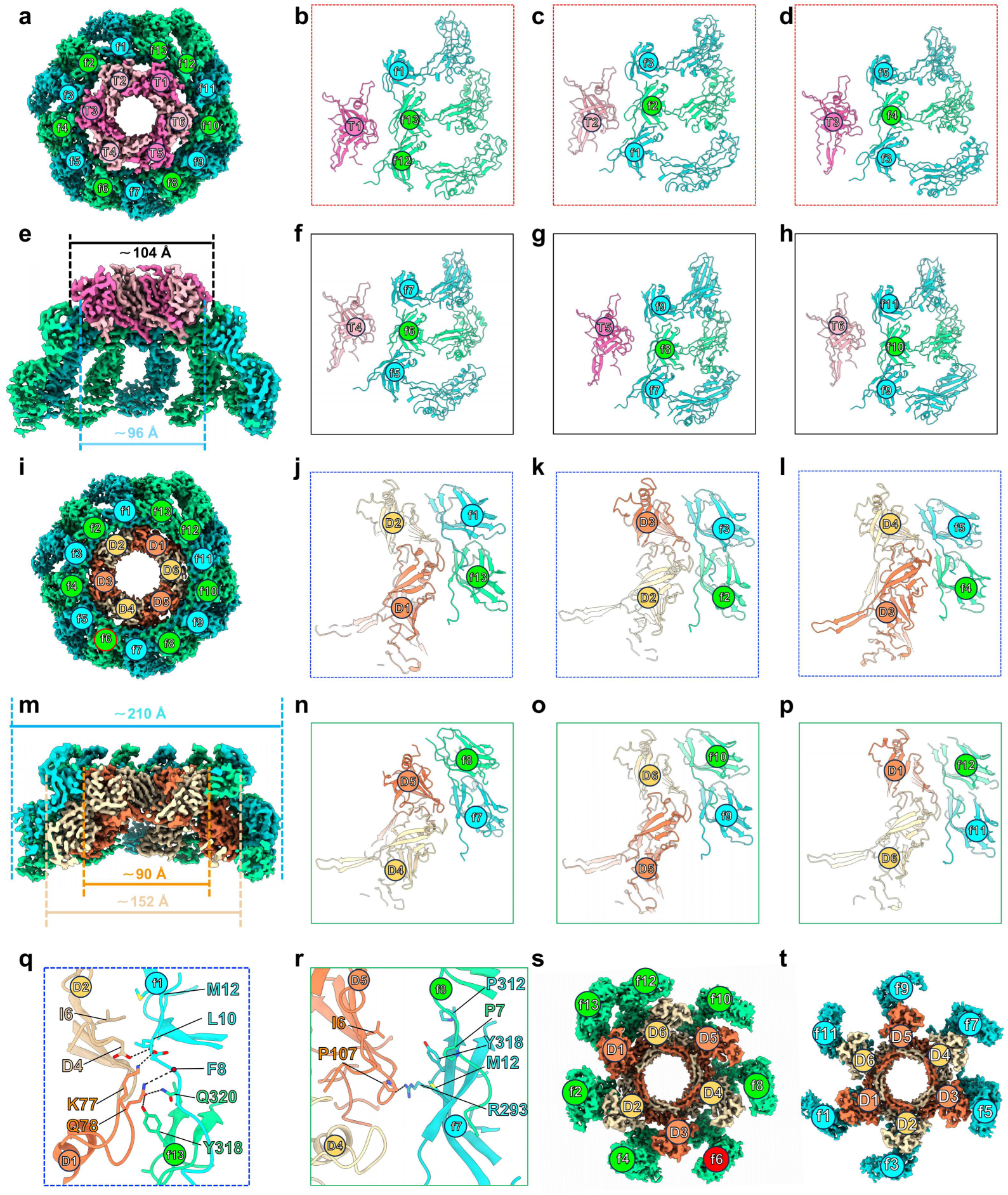
Symmetry-mismatched organization of the 13-subunit gp6 collar surrounding the C6-symmetric gp23-gp27 core. **a**, Top view of the cryo-EM density map showing the gp23 hexamer positioned above the 13-subunit gp6 collar. gp23 and gp6 subunits are labeled T1-T6 and f1-f13, respectively. **b-d**, Local arrangements of gp23 subunits T1-T3 and neighboring gp6 subunits. **e**, Side view of the gp23-gp6 assembly, showing the tilted arrangement of the collar relative to gp23. The gp23 ring and the central opening of the upper gp6 ring measure approximately 104 and 96 Å in diameter, respectively. **f-h**, Local arrangements of gp23 subunits T4-T6 and neighboring gp6 subunits, showing greater separation than in b-d. No specific gp23-gp6 interactions are resolved. **i**, Top view of the cryo-EM density map showing the gp27 hexamer embedded within the gp6 collar. gp27 subunits are labeled D1-D6. **j-l**, Three closely apposed interfaces between adjacent gp27 subunits and paired gp6 subunits: D1-D2 with f13-f1 (**j**), D2-D3 with f2-f3 (**k**), and D3-D4 with f4-f5 (**l**). **m**, Side view of the gp27-gp6 assembly. The upper and expanded lower regions of gp27 measure approximately 90 and 152 Å across, respectively, whereas the surrounding gp6 collar extends to approximately 210 Å. **n-p**, Three more widely separated interfaces: D4-D5 with f7-f8 (**n**), D5-D6 with f9-f10 (**o**), and D6-D1 with f11-f12 (**p**). **q-r**, Representative atomic details of the interfaces shown in j and n, respectively. **s-t**, Bottom views showing the lower-region arrangement of selected gp6 subunits relative to gp27. gp6 subunits f2, f4, f6, f8, f10, and f12 occupy spaces between adjacent gp27 subunits, with f13 positioned alongside D1 (**s**). gp6 subunits f1, f3, f5, f7, f9, and f11 lie beneath gp27 subunits D1-D6, respectively (**t**).

The gp27 hexamer is embedded within the 13-subunit gp6 collar (**Fig. 6i**), with its upper region closely apposed to the upper pseudo-C13 gp6 ring (**Fig. 6m**). Its expanded lower region is encircled by the lower tier of the gp6 collar (**Fig. 6m**). Viewed from the side, the upper gp6 ring surrounds the upper region of gp27 and lies beneath the outer rim of gp23, forming a nested arrangement at the gp23-gp27 junction (**Fig. 6i and 6m** ). Consistent with this organization, the upper gp6 ring measures approximately 96 Å in diameter (**Fig. 6e**), intermediate between the upper gp27 tier (approximately 90 Å) and the gp23 ring (approximately 104 Å) (**Fig. 6m**). Below this junction, gp27 expands to approximately 152 Å, while the surrounding gp6 collar extends outward to approximately 210 Å, forming the peripheral skirt of the baseplate (**Fig. 6m**).

The tilt of the gp6 collar also results in uneven spacing at the gp27-gp6 interface. In the upper region, each junction between adjacent gp27 subunits interacts with a corresponding junction between neighboring gp6 subunits (**Fig. 6i**). Specifically, the D1-D2, D2-D3, and D3-D4 junctions lie closer to their opposing gp6 junctions (**Fig. 6j-l**) and form hydrogen bonds and hydrophobic contacts (**Fig. 6q**). By comparison, the D4-D5, D5-D6, and D6-D1 junctions are farther from their opposing gp6 junctions (**Fig. 6n-p**), forming fewer hydrogen bonds and only weak hydrophobic contacts (**Fig. 6r**). The gp6 subunit f6 occupies a gap between these paired interfaces and shows no apparent contacts with the gp27 hexamer (**Fig. S11a**). Thus, the upper gp27-gp6 interface comprises six non-equivalent contact sites, with three forming more extensive interactions and the other three making limited contacts.

In the lower region, gp6 subunits f2, f4, f6, f8, f10, and f12 occupy the spaces between the expanded lower regions of adjacent gp27 subunits (**Fig. 6s**), without forming specific interactions. In contrast, gp6 subunits f1, f3, f5, f7, f9, and f11 lie directly beneath gp27 subunits D1–D6, respectively, while f13 lies alongside D1 (**Fig. 6t**). Despite this spatial proximity, detailed inspection reveals only potential electrostatic interactions, with most opposing surfaces lacking specific contacts (**Fig. S11b-d**).

Together, these observations suggest that the 6:13 symmetry mismatch is accommodated through a nested, asymmetric arrangement in which the gp6 collar contacts the gp23-gp27 core primarily through non-equivalent interfaces in the upper region of gp27. The absence of specific gp23-gp6 interactions, the unequal extent of the six upper gp27-gp6 interfaces, and the limited contacts in the lower region allow the 13-subunit collar to surround the conserved sixfold-symmetric core without requiring equivalent interactions or one-to-one subunit correspondence.

### Structural conservation and diversification of host-recognition modules in the D29 baseplate

To elucidate the molecular basis of host recognition by phage D29, we compared the corresponding baseplate proteins of D29, Bxb1^13^, and Douge^16^ to identify their conserved and divergent structural features. Structural superposition further revealed a highly conserved core among the DTPs. D29 gp27 superimposes closely with Douge gp17 and Bxb1 gp23, with Cα RMSD values of 1.43 and 0.83 Å, respectively (**Fig. 7a**). However, unlike D29 gp27 and Douge gp17, Bxb1 gp23 contains two additional peripheral CTE-like domains 1 and 2 (**Fig. 7a-c**). In Bxb1, these peripheral domains participate in a host-binding-associated conformational transition that expands the baseplate diameter from approximately 11 to 20 nm, thereby increasing the potential contact area with the bacterial cell surface^13^.

**Fig. 7.**
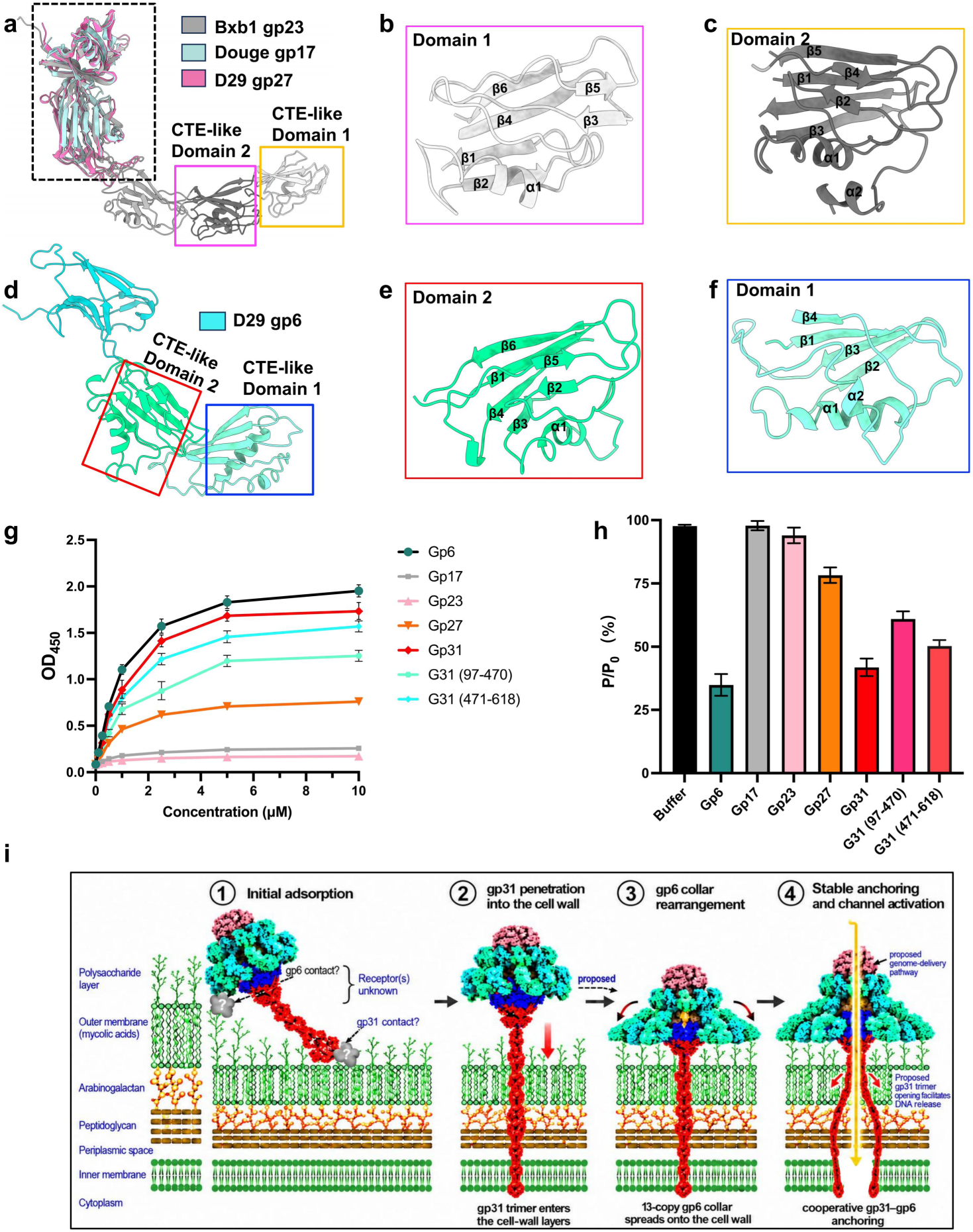
Host-cell binding activities of D29 baseplate proteins and proposed model of host engagement. **a**, Structural superposition of D29 gp27, Bxb1 gp23 and Douge gp17. **b-c**, Cartoon representations of CTE-like domains 1 (**b**) and 2 (**c**) of Bxb1 gp23. **d**, Overall structure of D29 gp6, showing its two CTE-like domains. **e-f**, Cartoon representations of gp6 domains 2 (**e**) and 1 (**f**), respectively. **g**, Concentration-dependent binding of the indicated D29 proteins to Msm cells, measured by ELISA at 450 nm. **h**, Protein-competition plaque assays showing the effects of the indicated proteins on D29 infectivity. Plaque numbers are expressed relative to the buffer control (*P/P₀* ). Data in g and h are presented as the mean ± s.d. of three independent experiments. **i**, Schematic representation of the proposed model of D29 baseplate-mediated host engagement.

Notably, the two peripheral CTE-like domains of Bxb1 gp23 exhibit clear structural similarity to those of D29 gp6 (**Fig. 7a-c**), although their domain assignments are reversed (**Fig. 7d-f**). CTE-like domain 1 of Bxb1 gp23 resembles CTE-like domain 2 of D29 gp6, with both containing a six-stranded β-sheet-rich core and a short α-helix (**Fig. 7b and 7e**). Conversely, CTE-like domain 2 of Bxb1 gp23 is structurally similar to CTE-like domain 1 of D29 gp6; both adopt compact α/β folds comprising a central β-sheet associated with two short α-helices (**Fig. 7c and 7f**). These similarities indicate that the peripheral domains of Bxb1 gp23 and D29 gp6 share a conserved modular structural framework. Their substantial sequence divergence, however, may generate distinct surface properties and interaction profiles, potentially contributing to differences in host recognition (**Fig. S12**).

In addition to the MTP gp6 collar, CFP gp31 forms a central fiber that projects from the distal end of the baseplate. Cryo-EM micrographs revealed substantial variation in its apparent length, measuring approximately 220 Å in most particles and extending to approximately 290 Å in a small subset (**Fig. S13a-b**). Although full-length gp31 comprises 618 residues, only its N-terminal residues 1-240 could be modeled into the C3-reconstructed cryo-EM density, albeit at relatively low local resolution, whereas the remaining C-terminal region was unresolved (**Fig. 5a-b**). These observations suggest considerable conformational heterogeneity of gp31, particularly in its distal region. AlphaFold3-based structural prediction suggests that gp31 comprises an N-terminal arm (residues 1-46), an N-terminal domain (residues 47-96), an elongated stalk-like region (residues 97-240), a distal tip domain (residues 241-470), and a C-terminal domain (CTD; residues 471-618) with a predicted folded structure (**Fig. S13c**). The N-terminal region anchors gp31 to the baseplate, while the stalk-like region extends along the central fiber axis, connecting the proximal attachment region to the unresolved distal domains. The absence of well-resolved cryo-EM density for the predicted distal domains suggests conformational flexibility that may facilitate host recognition and attachment.

An analogous trimeric central fiber was found in the T5-like phage DT57C^52^, where CFP pb4 forms a flexible, β-strand-rich axial fiber whose proximal region was resolved, closely resembling D29 gp31. Mycobacteriophage Douge likewise possesses a trimeric CFP, gp20^52^, implicated in initial host-surface engagement. Moreover, central-fiber-associated receptor-binding proteins in DT57C^52^, T5^53^, and λ^53^ recognize cell surface proteins BtuB, FhuA, and LamB, respectively, linking host recognition to infection triggering. These precedents support a potential role for D29 gp31 in exploratory host-surface engagement, although its cognate receptor and involvement in irreversible adsorption remain unknown.

### Proteins in baseplate involved in host binding

To assess the host-interacting potential of the structurally exposed baseplate proteins, we expressed and purified recombinant gp6, gp17, gp23, gp27, full-length gp31, and two gp31 fragments spanning residues 97-470 and 471-618. In whole-cell ELISA assays^54,55^, gp6 and full-length gp31 exhibited the strongest concentration-dependent binding to Msm, followed by gp31(471-618), gp31(97-470), and gp27, whereas gp17 and gp23 produced only weak signals close to background levels (**Fig. 7g**). The cell-binding activity of D29 gp6 is consistent with that reported for the corresponding minor tail protein gp6 of the closely related mycobacteriophage L5, which binds intact mycobacterial cells and has been developed as a mycobacterial capture reagent^27^.

To determine whether the observed cell binding could interfere with productive infection, we next performed protein-competition plaque assays^56,57^. Host cells were preincubated with 10 μM of each purified protein, and plaquing efficiency was normalized to that of the buffer-treated control and expressed as *P/P₀*^58^. gp6 and full-length gp31 produced the strongest inhibition, reducing *P/P₀* to approximately 35-40%. The two gp31 fragments also inhibited plaque formation, with gp31(471-618) and gp31(97-470) reducing *P/P₀* to approximately 50% and 60%, respectively. gp27 produced a weaker effect, with a *P/P₀* value of approximately 80%, whereas gp17 and gp23 had little or no detectable effect (**Fig. 7h**). This concordance between whole-cell binding and inhibition of infection resembles the paired ELISA and protein-competition results reported for other phage receptor-binding proteins, including the trimeric tail-fiber protein Pts_HN10M^54^, whose isolated C-terminal region retained bacterial cell-binding activity.

## Discussion

D29 is one of the most widely used subcluster A2 mycobacteriophages. Here, we determined cryo-EM structures of its capsid, connector, tail tube, and baseplate, revealing a near-complete virion architecture. These structures reveal two previously unidentified features at symmetry-mismatched interfaces. At the portal-capsid junction, five copies of the small accessory protein gp19.1 bridge neighboring capsid subunits around the dodecameric portal, providing local reinforcement at the fivefold vertex. At the baseplate, a 13-subunit gp6 collar surrounds the sixfold-symmetric gp23-gp27 core, with this unusual 6:13 mismatch accommodated primarily through non-equivalent gp27-gp6 contacts. D29 accommodates these symmetry mismatches in two different ways. Functional assays further implicate gp6 and the flexible central fiber gp31 in host binding, linking the distal architecture to candidate adsorption functions. Together, these findings establish a structural framework for investigating assembly and infection initiation in D29 and related A2 mycobacteriophages.

This study provides the first high-resolution structural description of a subcluster A2 mycobacteriophage. D29 shares a conserved virion framework with Bxb1^13^ and Douge^16^, comprising an HK97-fold capsid, a portal-containing connector, and a long non-contractile tail. Its capsid and tail structures closely resemble those of Bxb1^13^, whereas its overall baseplate organization is more similar to that of Douge^16^. Despite these similarities, D29 lacks the capsid and tail-tube extensions and elaborate tail-tip cage described in Bxb1^13^, as well as the widely distributed

Ig-like glycan-binding domains found in Douge^16^. D29 instead features a gp19.1-reinforced portal vertex and a distinctive 13-subunit gp6 collar surrounding the conserved sixfold-symmetric baseplate core. Notably, the peripheral domains of gp6 structurally resemble domains incorporated into the Bxb1 distal tail protein gp23, although they occupy different positions within their respective assemblies. The approximately circular D29 gp6 collar is ∼210 Å in diameter, giving it a footprint comparable to that of the host-bound, expanded Bxb1 gp23 assembly (∼200 Å). The D29 gp6 collar may expand further upon host binding, potentially providing an even larger contact surface^13^. This similarity suggests that distinct baseplate architectures can provide comparable potential host-contacting areas, although host-induced expansion of the D29 collar remains unverified. Together, these comparisons reveal how related structural modules can be arranged differently within a conserved virion framework, expanding our understanding of mycobacteriophage structural diversity.

Previous studies have not established which proteins directly recognize host receptors in D29-like phages. However, earlier studies showed that the minor tail protein gp6 of mycobacteriophage L5 binds mycobacterial cells and identified a homolog (gp6) in D29^27,28^. Our structures show that D29 gp6 is located at the exposed periphery of the baseplate and shares structural similarities with the host-interacting peripheral domains of Bxb1 gp23^13^. Together with its whole-cell binding activity and inhibition of plaque formation in competition assays, these findings strongly support gp6 as an important host-binding component. The central fiber gp31 occupies a position similar to that of Douge gp20 and likewise adopts a trimeric organization^16^. Its cell-binding activity and inhibition of plaque formation further support its involvement in host engagement. Together, these structural and functional results identify gp6 and gp31 as components of the D29 host-binding apparatus, suggesting that both the peripheral collar and the central fiber contribute to adsorption. These findings provide a basis for defining their respective contributions to attachment and identifying their cognate host receptors.

Specific surface glycolipids are important determinants of mycobacteriophage host range. BPs and Muddy^59^, for example, depend on trehalose polyphleates (TPPs) for efficient adsorption and infection, and bacteria lacking these glycolipids become resistant. However, a single amino acid substitution in their respective tail spike proteins can enable infection in the absence of TPPs^59^. This precedent highlights the D29 adsorption apparatus, particularly gp6 and gp31, as a potential target for structure-guided engineering. Understanding how these components cooperate during infection will be essential for such efforts.

Our structural and functional findings support a tentative multistep model for D29 adsorption and infection initiation (**Fig. 7i**). Either gp31 or gp6 may initiate contact with the mycobacterial surface. The flexible gp31 trimer could explore the surface through its distal C-terminal region, while the gp6 collar may provide multivalent contacts that stabilize attachment. Further engagement could bring the baseplate closer to the cell envelope, potentially allowing gp31 to penetrate its outer layers and the gp6 collar to rearrange or spread outward. Additional components associated with the unresolved distal region of gp31 might also contribute to binding or local envelope degradation, although their presence remains unconfirmed. These cooperative interactions could promote conformational changes in the central baseplate, leading to distal-channel opening and genome delivery. The receptors, sequence of events, and proposed structural transitions remain to be established, but this model provides a testable framework for D29 infection initiation.

Overall, this study reveals how D29 combines a gp19.1-reinforced portal vertex with a symmetry-mismatched baseplate organized around a 13-subunit gp6 collar. Structural and functional analyses further identify gp6 and gp31 as host-binding components, linking the baseplate architecture to adsorption. These findings advance our understanding of mycobacteriophage assembly and host interaction and provide a foundation for future studies of receptor recognition and infection initiation.

## Methods

### Bacterial culture and phage propagation

*Mycobacterium smegmatis* mc²155 was used for the propagation and functional characterization of mycobacteriophage D29. Bacteria were cultured at 37°C in Middlebrook 7H9 broth supplemented with 10% albumin-dextrose-catalase (ADC), 0.5% glycerol, 0.05% Tween 80, and 1 mM CaCl₂. For growth on solid medium, Middlebrook 7H10 agar supplemented with 10% ADC and 0.5% glycerol was used.

An initial D29 stock was prepared using the double-layer agar method. Briefly, exponentially growing M. smegmatis cells were mixed with D29 and molten Middlebrook top agar containing 1 mM CaCl₂. The mixture was poured onto 7H10 agar plates and incubated at 37°C until confluent lysis was observed. Phage particles were recovered by overlaying the plates with SM buffer (50 mM Tris–HCl, pH 7.5, 70 mM NaCl, 8 mM MgSO₄, and 0.01% gelatin). The resulting phage suspension was clarified by centrifugation and passed through a 0.22 μm membrane filter.

For large-scale amplification, the plate-derived phage stock was used to infect an exponentially growing suspension of Msm. Following incubation until extensive bacterial lysis was observed, the culture was centrifuged to remove cells and cellular debris, and the phage-containing supernatant was collected. Residual host nucleic acids were digested with DNase I and RNase A for 30 min at room temperature. The clarified lysate was then centrifuged at 30,000 × g for 2 h at 4°C. The resulting phage pellet was gently resuspended in SM buffer and loaded onto a 15-60% (w/v) sucrose density gradient. Following centrifugation at 45,000 × g for 3 h at 4°C, the visible phage-containing band was collected and dialyzed overnight against SM buffer to remove sucrose. The purified D29 particles were subsequently concentrated by ultrafiltration and stored at 4°C until cryo-EM grid preparation.

### Cryo-EM grid preparation and data collection

The 4 μL aliquots of purified D29 particles were applied to freshly glow-discharged Quantifoil R2/1, 300-mesh gold grids. Grids were prepared at 4°C and 100% relative humidity using a Vitrobot Mark IV. After incubation for 10 s, excess liquid was removed by blotting for 3 s at a blot force of 4, and the grids were plunge-frozen in liquid ethane cooled by liquid nitrogen.

Cryo-EM data were collected using a 300-kV Titan Krios G4 microscope (Thermo Fisher Scientific) equipped with a Falcon 4 direct electron detector and a Selectris X energy filter set to a slit width of 10 eV. Movies were automatically acquired using EPU software (Thermo Fisher Scientific) at a nominal magnification of 105,000 ×, corresponding to a calibrated pixel size of 1.2 Å per pixel. Movies were recorded in electron-event representation (EER) mode, with an accumulated total dose of approximately 50 e⁻/Å². The EER fractions were grouped during motion correction. The nominal defocus range was set between -0.8 and -2.4 μm. In total, 7585 movies were collected for subsequent image processing.

### Cryo-EM single-particle analysis

A total of 7,585 motion-corrected micrographs were imported into cryoSPARC v4.6.2^60^. Contrast transfer function (CTF) parameters were estimated using Patch CTF Estimation, and 7,469 high-quality micrographs were retained after manual curation using Manually Curate Exposures.

Capsid reconstruction and localized reconstruction of the tail root. For capsid reconstruction, an initial set of 11,134 particles was picked from 1,000 micrographs using Blob Picker and subjected to reference-free 2D classification to generate templates. Subsequent template-based picking across the complete dataset yielded 104,003 particles. After 2D classification and particle selection, 56,131 particles were subjected to Ab-Initio Reconstruction and Heterogeneous Refinement with icosahedral (I) symmetry imposed, yielding a well-resolved class containing 55,842 particles. These particles were further processed using Global CTF Refinement, Local CTF Refinement, and Non-Uniform (NU) Refinement with I symmetry, resulting in a reconstruction at an overall resolution of 2.49 Å.

To resolve the capsid-attached tail root, the particle stack obtained before Ewald sphere correction was symmetry-expanded according to icosahedral symmetry. The symmetry-expanded particles were subjected to volume alignment and focused 3D classification without imposed symmetry, using a mask centered on a 5-fold vertex, to isolate the class containing the tail root. After duplicate removal, 58,989 symmetry-expanded subparticles were retained. Owing to the limitations of focused classification, this dataset included a small number of false-positive subparticles derived from tailless capsids. Independent NU refinements with C5 and C6 symmetry imposed yielded maps of the C5-symmetric outer vertex region and the C6-symmetric tail root at overall resolutions of 2.92 Å and 2.81 Å, respectively.

Tail shaft reconstruction. For reconstruction of the repetitive tail shaft, 352,270 candidate particles were initially picked from 1,000 micrographs using Blob Picker. Representative 2D class averages derived from 43,167 selected particles were subsequently used as templates for Filament Tracer, which identified 846,010 particles across the complete dataset. After 2D classification and removal of low-quality particles, 692,602 particles were retained and subjected to Ab-Initio Reconstruction and Heterogeneous Refinement. Three well-resolved classes comprising a total of 565,700 particles were pooled and subjected to NU Refinement followed by Local Refinement. Because of the pronounced and irregular bending of the flexible tail, the final local refinement was performed using a single-particle approach without imposing helical symmetry, yielding a tail-shaft map at an overall resolution of 3.20 Å.

To reconstruct the distal tail tip, 1,995 particles were manually picked from 180 micrographs and subjected to 2D classification. Four representative tail-tip classes were selected as templates for template-based picking, yielding 962,415 candidate particles. After 2D classification and particle selection, 42,098 particles were subjected to ab initio reconstruction and heterogeneous refinement. A class containing 36,229 particles was selected and refined to 3.85 Å using non-uniform (NU) refinement. These particles were then used to train a dedicated Topaz model for particle picking across the complete dataset. Further 2D classification, ab initio reconstruction, and heterogeneous refinement of the Topaz-picked particles yielded a final stack of 42,979 particles.

Initial NU refinement without imposed symmetry (C1) produced a 3.81 Å reconstruction. Global and local CTF refinement, followed by C3-symmetric NU refinement, yielded a 2.88 Å map with improved definition of the central body. To better resolve the flexible peripheral and distal regions, the particles were aligned to the distal tip using volume alignment and re-extracted. The re-extracted particles were processed through two complementary refinement branches. In the C1 branch, NU refinement yielded a 3.24 Å reconstruction, and subsequent 3DFlex refinement produced a 3.32 Å map with improved continuity of the peripheral density. Map enhancement further clarified the 13-copy organization of the gp6 collar. In the C3 branch, NU refinement yielded a 3.01 Å reconstruction, followed by 3DFlex refinement to 3.16 Å. This branch improved the continuity of the distal tail-tip density, with subsequent map enhancement more clearly delineating the resolved trimeric region of the gp31 central fiber.

Empty capsid and associated tail root. During 2D classification of the 104,003 template-picked capsid particles, a subpopulation of 2,874 particles corresponding to empty capsids was identified. These particles were subjected to Ab-Initio Reconstruction with I symmetry, followed by Heterogeneous Refinement without imposed symmetry (C1), yielding a class containing 2,607 particles. Subsequent I-symmetric NU Refinement produced an empty-capsid reconstruction at an overall resolution of 4.22 Å. Following icosahedral symmetry expansion, volume alignment, focused 3D classification, and duplicate removal, 2,587 symmetry-expanded subparticles were retained. NU Refinement with C6 symmetry yielded a reconstruction of the tail root associated with the empty capsid at an overall resolution of 4.04 Å.

Resolution estimation and map postprocessing. All cryo-EM single-particle image-processing steps-including CTF refinement, symmetry expansion, 3D classification, NU Refinement, 3DFlex Refinement, and local-resolution estimation-were performed in cryoSPARC v4.6.2. All reported resolutions were determined from gold-standard Fourier shell correlation curves using the FSC = 0.143 criterion and solvent-corrected masks. Selected final density maps were enhanced using DeepEMhancer in highRes mode, with the unfiltered half-maps supplied as input. For maps obtained by 3DFlex Refinement, the solvent mask generated during 3DFlex Refinement was supplied as a binary mask to minimize boundary artifacts. Density maps and local-resolution surface representations were visualized and rendered using UCSF ChimeraX^61^.

### Model building and refinement

Atomic models of the D29 capsid, portal-capsid interface, connector, tail tube, and baseplate were constructed using a common workflow. Initial models of the individual D29 structural proteins were generated using AlphaFold3 (AF3) and used as starting templates. The predicted models were rigid-body fitted into the corresponding cryo-EM densities using the “Fit in Map” function in UCSF Chimera^62^ and ChimeraX^61^.

For the capsid, the AF3-predicted gp17 model was initially fitted into the icosahedrally reconstructed map and subsequently adjusted to match the local density. Symmetry-related copies were generated to construct the complete capsid model. The gp17 subunits surrounding the portal vertex were independently fitted and refined against the symmetry-free localized reconstruction to account for their distinct conformations. The gp19.1 model was placed into the additional density at the portal-capsid interface, and its sequence register and interactions with the surrounding gp17 subunits were manually adjusted according to the resolved backbone and side-chain features.

For the connector and tail tube, the predicted models of the corresponding component proteins were individually docked into their locally reconstructed maps. Symmetry-related copies were generated according to the oligomeric organization of each ring, followed by manual adjustment of intersubunit interfaces. The portal and adaptor models were refined according to their 12-fold organization, whereas the stopper, terminator, and tail-tube components were modeled according to their 6-fold organization.

For the baseplate, AF3-predicted models of gp6, gp23, ,gp26, gp27, gp28, and the resolved region of gp31 were separately fitted into the corresponding densities. Because the baseplate contains symmetry-mismatched C6, C13, and C3 assemblies, each component was fitted into the symmetry-free localized reconstruction without imposing a common symmetry constraint. Individual subunits were subsequently adjusted to accommodate local conformational differences and the heterogeneous gp23-gp6 and gp27-gp6 interfaces. Only regions supported by interpretable cryo-EM density were included in the final atomic models.

All models were manually inspected and rebuilt in COOT^63^, including correction of backbone geometry, sequence registration, side-chain conformations, and flexible regions. Multiple rounds of real-space refinement were performed using the cryo-EM module in PHENIX^64^, supplemented by iterative refinement with Namdinator^65^ to improve model-map agreement and stereochemical quality. Secondary-structure, Ramachandran, and noncrystallographic-symmetry restraints were applied where appropriate. Final models were assessed using the comprehensive cryo-EM validation tools implemented in PHENIX^64^, including model geometry, steric clashes, and model-to-map agreement. Structural figures, surface representations, and domain-level illustrations were generated using UCSF Chimera^62^, ChimeraX^61^, and PyMOL (https://pymol.org/2/).

### Cloning, expression, and purification of gp6, gp17, gp23, gp27, gp31, gp31(97-470), and gp31(471-618)

The coding sequences of D29 gp6, gp17, gp23, gp27, and gp31, together with two gp31 fragments comprising residues 97-470 and 471-618, were amplified from the D29 genome by PCR. The amplified fragments were inserted into a modified pET-28a expression vector containing an N-terminal His₆ tag using [restriction digestion and ligation/Gibson assembly]. All constructs were verified by DNA sequencing and transformed into *Escherichia coli* BL21(DE3) cells.

Transformed cells were cultured at 37°C in LB medium containing [50 μg/mL kanamycin or 100 μg/mL ampicillin]. When the cultures reached an OD₆₀₀ of approximately 0.8, protein expression was induced with 0.5 mM IPTG, followed by incubation at 16°C for 18 h. Cells were collected by centrifugation and resuspended in lysis buffer containing 50 mM Tris-HCl, pH 8.0, 500 mM NaCl, 20 mM imidazole, 5% glycerol, and 1 mM DTT. After disruption by sonication, cell debris was removed by centrifugation at 20,000 × g for 1 h at 4°C.

The clarified supernatant was loaded onto a Ni-affinity column pre-equilibrated with lysis buffer. The column was washed with buffer containing 60 mM imidazole, and bound proteins were eluted with 300 mM imidazole. Protein-containing fractions were pooled, concentrated, and further purified by size-exclusion chromatography using a Superdex 200 Increase 10/300 GL column equilibrated with 20 mM HEPES, pH 7.5, 150 mM NaCl, and 1 mM DTT. Peak fractions were analyzed by SDS-PAGE, pooled, and concentrated. Purified proteins were used immediately for bacterial binding and plaque-formation assays or flash-frozen in liquid nitrogen and stored at -80°C.

### Bacterial binding by whole-cell ELISA

The binding of recombinant D29 structural proteins to Mycobacterium smegmatis mc²155 cells was evaluated using a whole-cell enzyme-linked immunosorbent assay (ELISA). M. smegmatis cells were grown in Middlebrook 7H9 medium supplemented with 10% ADC, 0.5% glycerol, 0.05% Tween 80, and 1 mM CaCl₂ at 37°C to the mid-logarithmic phase (OD₆₀₀ = 0.6-0.8). Cells were harvested by centrifugation at 5,000 × g for 5 min, washed three times with phosphate-buffered saline (PBS) to remove residual culture medium and Tween 80, and resuspended in PBS to an OD₆₀₀ of 0.2.

Aliquots of the bacterial suspension (100 μL per well) were added to high-binding 96-well microplates and incubated overnight at 4°C to allow cell immobilization. After removal of the suspension, the immobilized cells were fixed with 4% paraformaldehyde for 20 min at room temperature and washed three times with PBS containing 0.05% Tween 20 (PBST). Nonspecific binding sites were blocked with 3% skim milk in PBS for 1 h at 37°C.

Purified His-tagged gp6, gp17, gp23, gp27, full-length gp31, gp31(97-470), or gp31(471-618) was serially diluted in PBS and added to the wells at the indicated concentrations. After incubation for 1 h at 37°C, the wells were washed five times with PBST. Bound proteins were detected using an anti-His-tag primary antibody, followed by an HRP-conjugated secondary antibody. After five additional washes, 3,3′,5,5′-tetramethylbenzidine (TMB) substrate was added and the reaction was developed for 10 min at room temperature. The reaction was stopped with 2 M H₂SO₄, and absorbance was measured at 450 nm using a microplate reader.

Wells containing immobilized bacteria but no recombinant protein and wells without bacterial cells were included to determine background signals. An unrelated His-tagged protein or bovine serum albumin was used as an additional negative control where indicated. Background absorbance from wells lacking bacterial cells was subtracted from the corresponding measurements. Each condition was analyzed in triplicate, and the experiment was independently repeated three times.

### Protein-competition plaque assay

The effects of purified D29 structural proteins on phage infection were examined using a protein-competition plaque assay. *Mycobacterium smegmatis* mc²155 cells were grown to the mid-logarithmic phase, harvested by centrifugation, washed twice, and resuspended in Tween-free Middlebrook 7H9 medium supplemented with 1 mM CaCl₂ to approximately 1 × 10⁸ CFU mL⁻¹. Aliquots of the bacterial suspension (100 μL) were mixed with purified gp6, gp17, gp23, gp27, full-length gp31, gp31(97-470), or gp31(471-618) at a final concentration of 10 μM, unless otherwise indicated. The mixtures were incubated at 37°C for 20 min to allow the recombinant proteins to bind to the bacterial surface.

Approximately 100-200 PFU of D29 were subsequently added to each mixture, followed by incubation at 37°C for 10 min. The samples were immediately mixed with 3 mL of molten Middlebrook top agar containing 0.7% agar and 1 mM CaCl₂ and poured onto Middlebrook 7H10 agar plates supplemented with 10% ADC and 0.5% glycerol. After solidification, the plates were incubated at 37°C for 24 h, and plaques were counted.

Samples containing an equivalent volume of protein storage buffer without recombinant protein were used as vehicle controls. Relative plaque-forming efficiency was expressed as *P/P_0_* and calculated as: *P/P_0_* = N_protein-treated_/N_buffer control_, where N_protein-treated_/ and N_buffer control_ represent the numbers of plaques obtained from the recombinant-protein-treated and buffer-control groups, respectively. Protein-treated bacterial cells without phage were plated in parallel to exclude direct effects of the recombinant proteins on bacterial growth or lawn formation. Each condition was tested in triplicate, and the experiment was independently repeated three times.

### Statistical analysis

Data are presented as the mean ± standard deviation from three independent experiments. Statistical significance between each recombinant protein and the buffer-treated control was evaluated using one-way analysis of variance followed by Dunnett’s multiple-comparison test. A value of P < 0.05 was considered statistically significant.

## Data Availability

Cryo-EM maps of this study have been deposited in the Electron Microscope Data Bank (EMDB) with the accession codes XX and XX. The atomic coordinates and structure factors for the structures determined in this study have been deposited in the Protein Data Bank under the accession codes XX and XX.

## Acknowledgements

This study was supported by the Nation Key R&D Program of China (2021YFA1300903 to X.X., 2021YFA1300904 to T.Z., 2022YFC2304800 to J.H.), National Natural Science Foundation of China (32501089 to B.Z., 82341085 to X.X. and 32201153 to X.Z.), China Postdoctoral Science Foundation (2025M772866 to B.Z.), Major Project of Guangzhou National Laboratory (GZNL2024A01030 to J.H.), Guangzhou Municipal Science and Technology Program-Key R&D Initiatives (2024B03J1345 to J.H.), The Key Medical Disciplines and Specialties Program of Guangzhou (Tuberculosis) 2025-2027 (L.C.). The Science and Technology Planning Project of Fujian Province (2023J01292 to X.Z.). We thank the staffs at cryo-EM Facilities of GIBH-CAS, Guangzhou National Laboratory Bio-imaging Technology Platform for help on cryo-EM sample preparation and data collection.

## Contributions

B.Z., T.Z., L.C., X.Z. and X.X. conceived the study; B.Y. propagated the phage and performed the plaque assays; D.H. and Z.H. expressed and purified the recombinant proteins; X.Z. and J.G. performed the whole-cell ELISA assays; M.L. carried out cryo-EM data collection; Z.L. processed the cryo-EM data; Y.Z. refined and corrected the atomic models; B.Z., X.X., L.W., and X.Z. analyzed the cryo-EM structures; B.Z. and X.X. prepared the figures and wrote the initial manuscript, which was edited by all authors; B.Z., T.Z., L.C., J.H., X.Z. and X.X. acquired funding and supervised the project.

---

**Supplementary Fig. 1.**
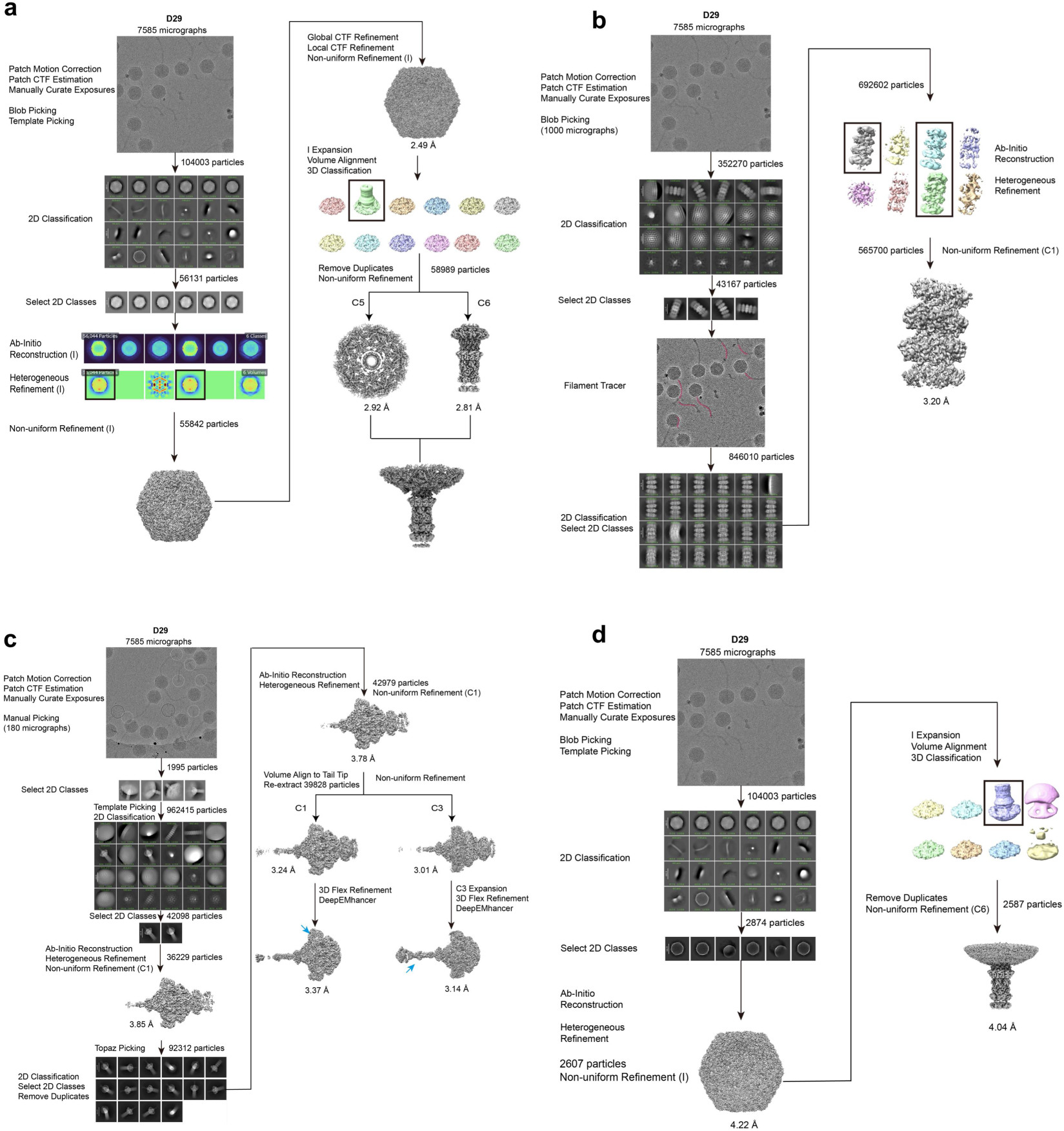
Cryo-EM data processing pipelines for the D29 structural reconstructions. **a**, Data-processing pipeline for the DNA-filled capsid and capsid-tail junction datasets. **b**, Data-processing pipeline for the tail tube dataset. **c**, Data-processing pipeline for the distal baseplate dataset. **d**, Data-processing pipeline for the genome-free capsid and associated tail-root datasets. Black boxes indicate selected classes, and black boxes indicate selected classes.

**Supplementary Fig. 2.**
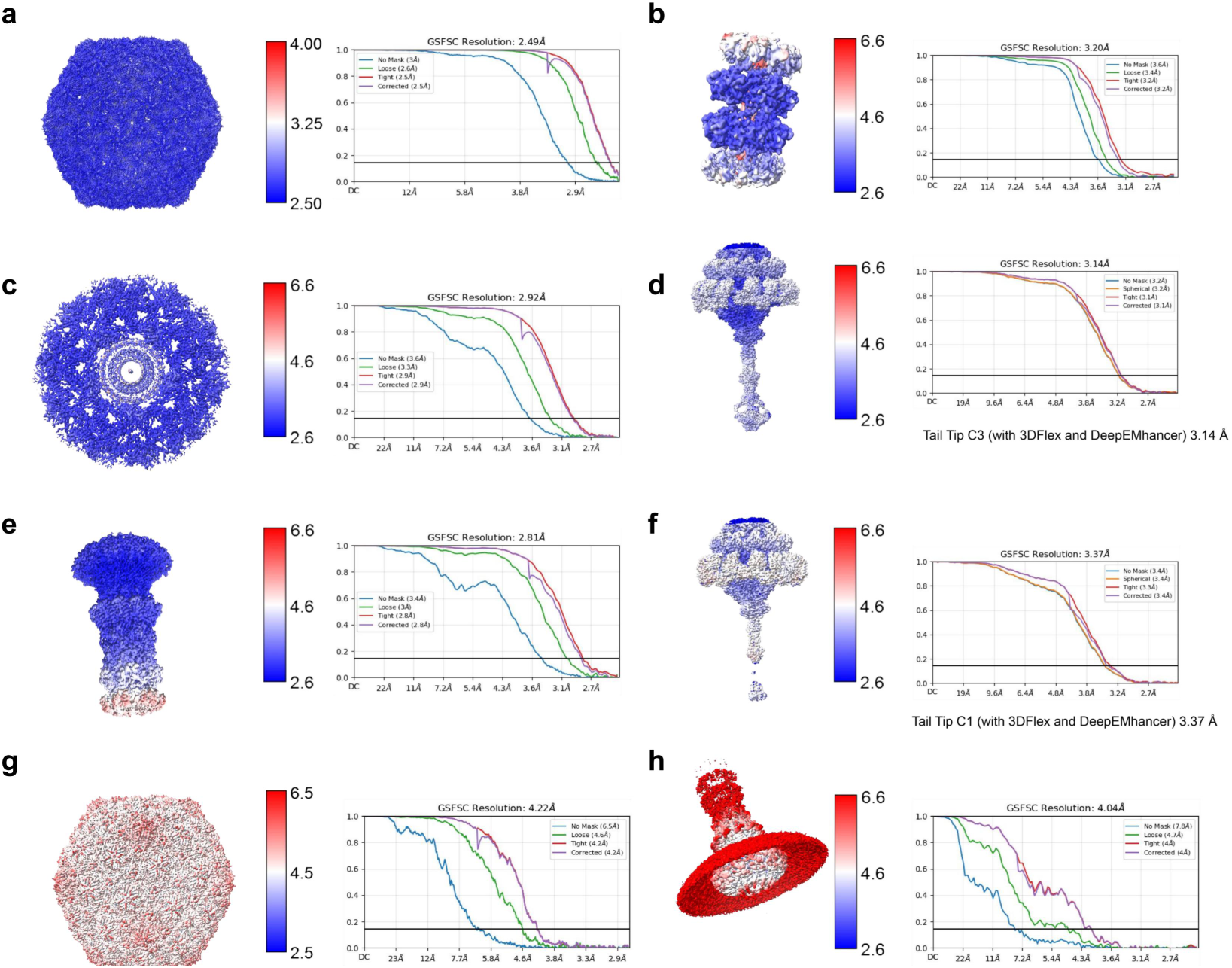
Resolution assessment of the D29 cryo-EM structures. **a-h**, Local- and global-resolution assessments of the DNA-filled capsid (**a**), tail tube (**b**), C5 capsid-portal interface (**c**), C3 tail tip (**d**), C6 connector (**e**), C1 tail tip (**f**), genome-free capsid (**g**) and genome-free connector (**h**) reconstructions, respectively. Global resolutions were determined by Fourier shell correlation (FSC) at the 0.143 criterion.

**Supplementary Fig. 3.**
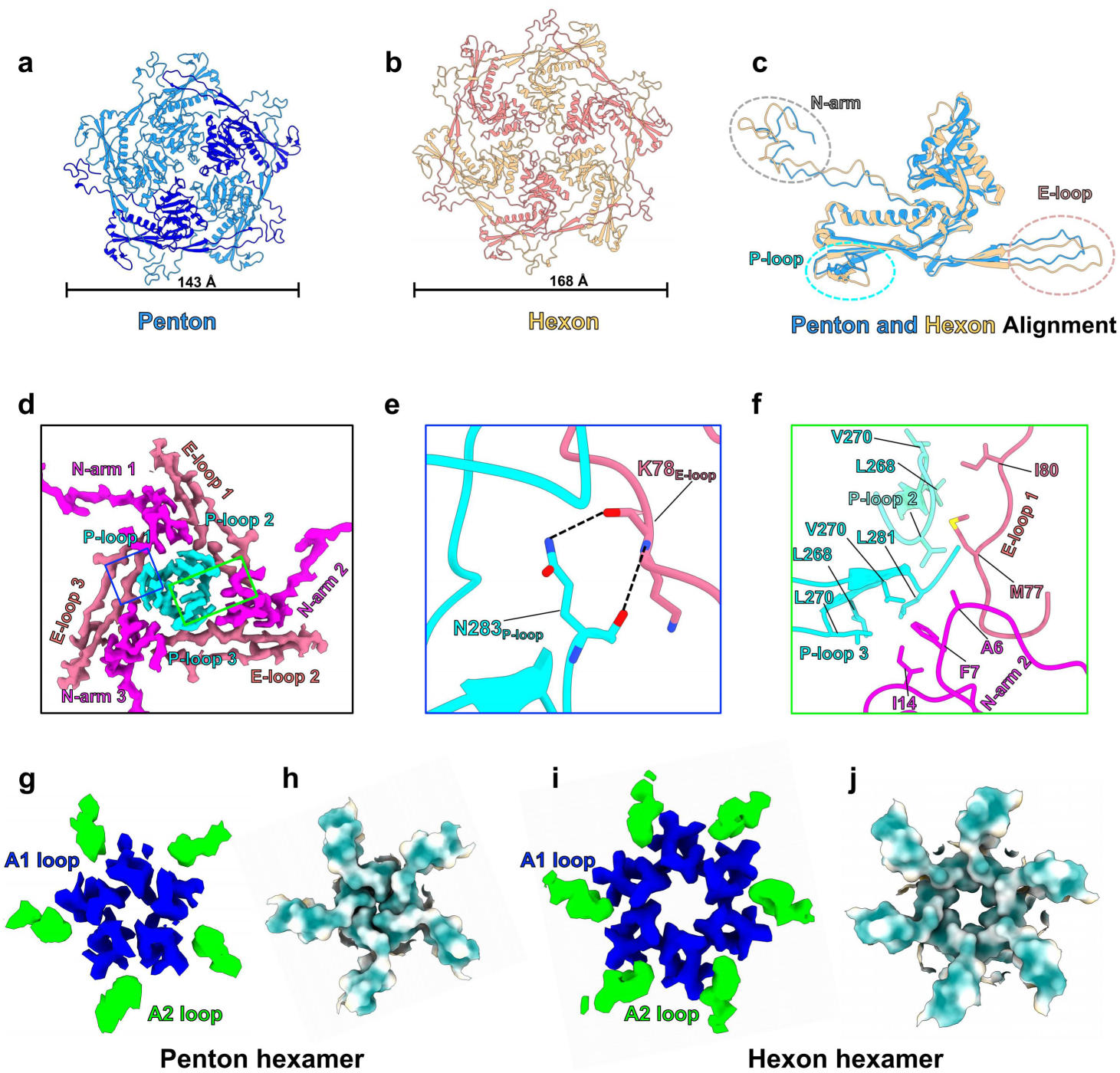
Structural organization and inter-subunit interactions of D29 gp17 capsomers. **a-b**, Cartoon representations of the gp17 penton (**a**) and hexon (**b**), with their dimensions indicated. **c**, Structural alignment of representative gp17 subunits from the penton and hexon, highlighting conformational differences in the N-arm, P-loop and E-loop. **d**, Cryo-EM map showing the interactions among the P-loops, E-loops and N-arms of three neighboring gp17 subunits at a capsid threefold axis. **e-f**, Cartoon representations of the boxed regions in **d**, showing the interaction between the E-loop and P-loop (**e**) and the interactions among the N-arm, E-loop and P-loop (**f**). Hydrogen bonds are indicated by dashed lines. **g-j**, Cryo-EM maps and hydrophobic surface representations of the A1 and A2 loops surrounding the penton (**g-h**) and hexon (**i-j**) axes.

**Supplementary Fig. 4.**
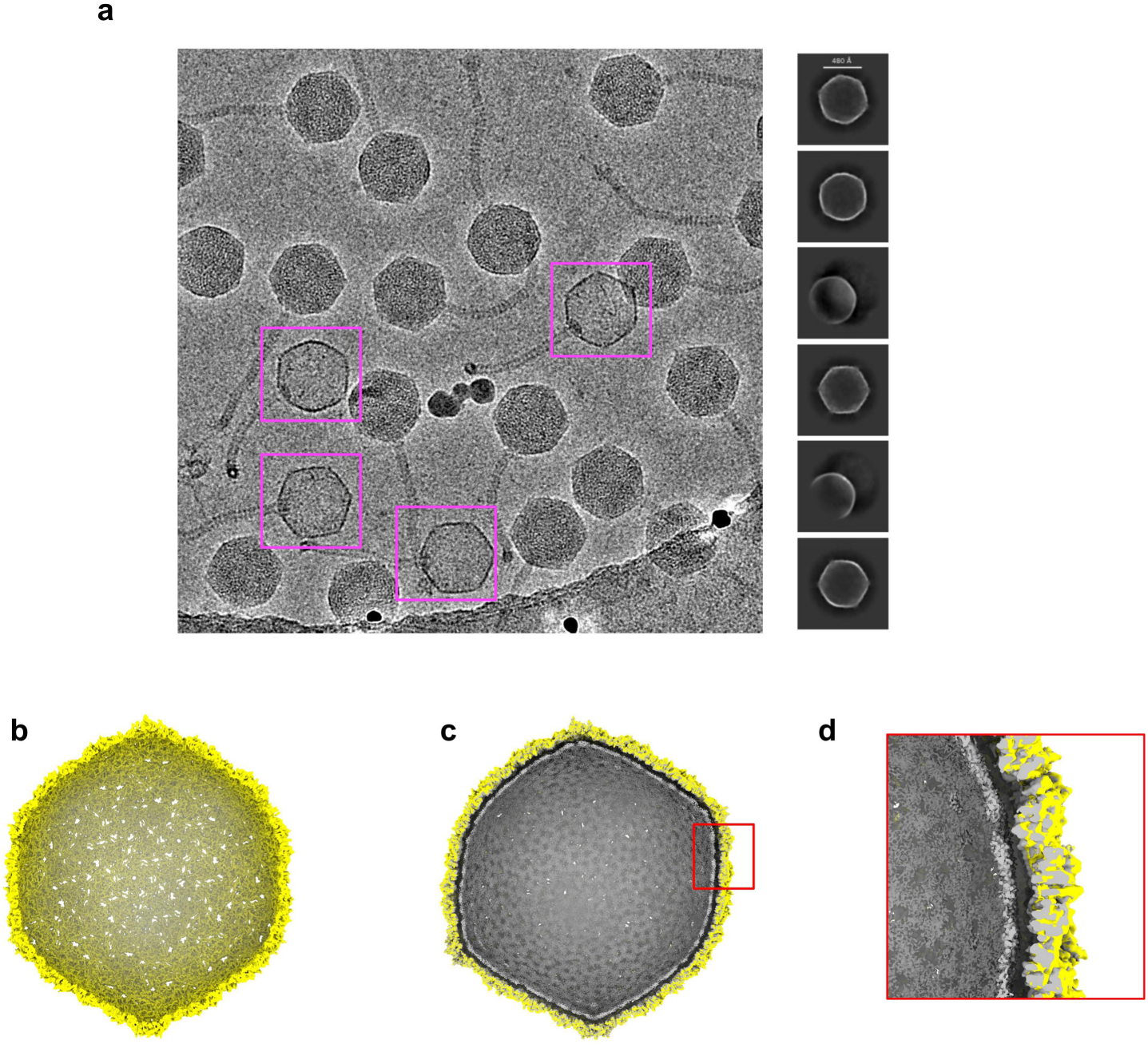
Structural comparison of genome-packed and genome-free D29 capsids. **a**, Representative cryo-EM micrograph and 2D class averages of genome-free D29 capsids. Genome-free particles are highlighted by magenta boxes. **b**, Cryo-EM reconstruction of the genome-free capsid, colored yellow. **c**, Cross-sectional view of the superimposed genome-packed and genome-free capsid reconstructions, colored gray and yellow, respectively. **d**, Enlarged view of the boxed region in **c**, highlighting the outward expansion of the genome-free capsid shell relative to the genome-packed capsid.

**Supplementary Fig. 5.**
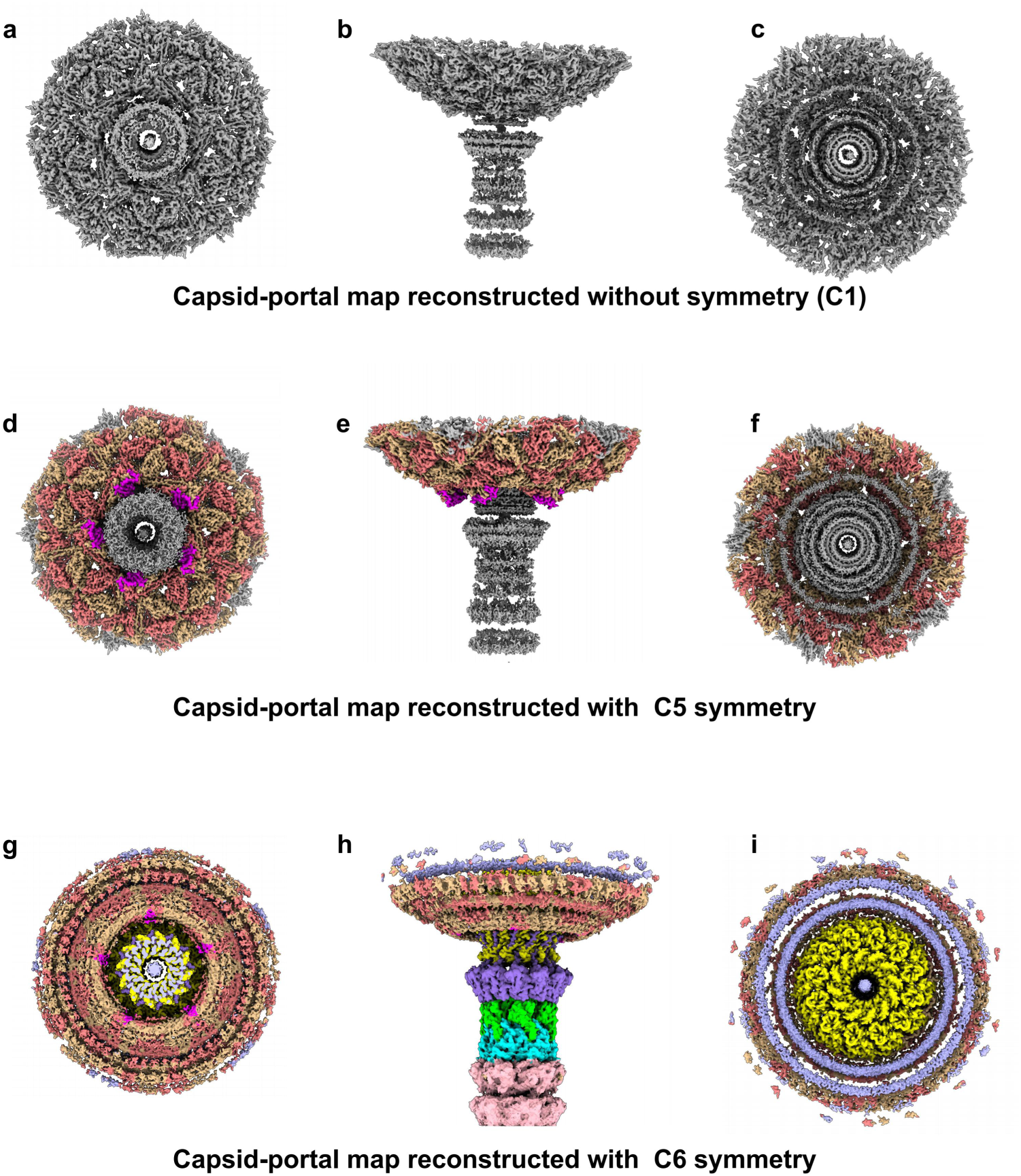
Cryo-EM reconstructions of the D29 capsid-portal junction with different imposed symmetries. **a-c**, Cryo-EM map of the capsid-portal junction reconstructed without imposed symmetry (C1), shown in two opposing axial views (a and c) and a side view (b). **d-f**, Cryo-EM map of the capsid-portal junction reconstructed with C5 symmetry, shown in two opposing axial views (d and f) and a side view (e). **g-i**, Cryo-EM map of the capsid-portal junction reconstructed with C6 symmetry, shown in two opposing axial views (g and i) and a side view (h).

**Supplementary Fig. 6.**
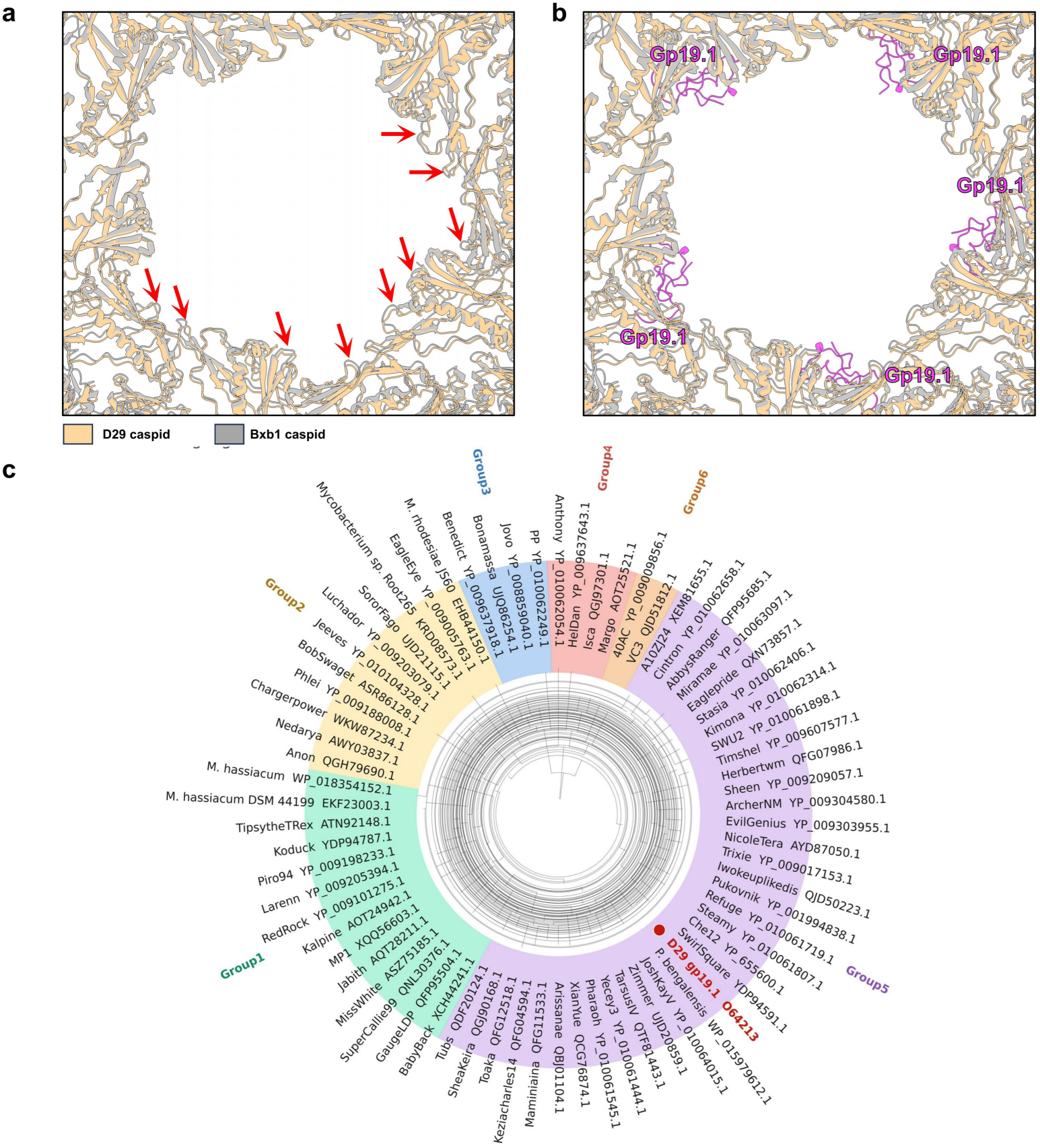
Structural comparison and phylogenetic distribution of gp19.1. **a**, Cartoon representations of the superimposed capsid subunits surrounding the portal vertices of D29 and Bxb1, colored tan and gray, respectively. Red arrows indicate the enlarged inter-subunit gaps in the D29 portal-proximal capsid lattice. **b**, Cartoon representation showing five gp19.1 molecules positioned within these gaps at the D29 portal-capsid junction. Gp19.1 is colored magenta. **c**, Phylogenetic tree of putative gp19.1 homologs identified in diverse mycobacteriophages and mycobacteria. The homologs are classified into six groups, indicated by different background colors, with D29 gp19.1 highlighted in red.

**Supplementary Fig. 7.**
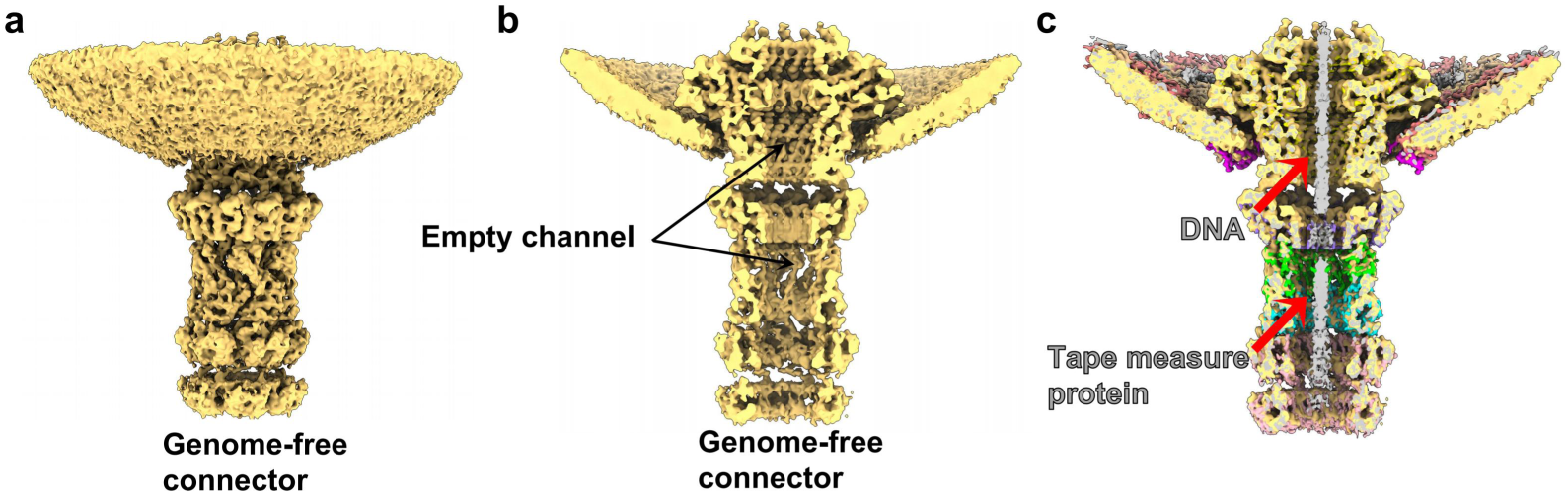
Structure of the genome-free D29 connector. **a**, Cryo-EM map of the connector associated with the genome-free capsid. **b**, Cutaway view showing the continuous density-free channel through the genome-free connector and proximal tail. **c**, Cutaway view of the genome-packed D29 connector for comparison, showing the axial densities corresponding to dsDNA and the tape measure protein gp26.

**Supplementary Fig. 8.**
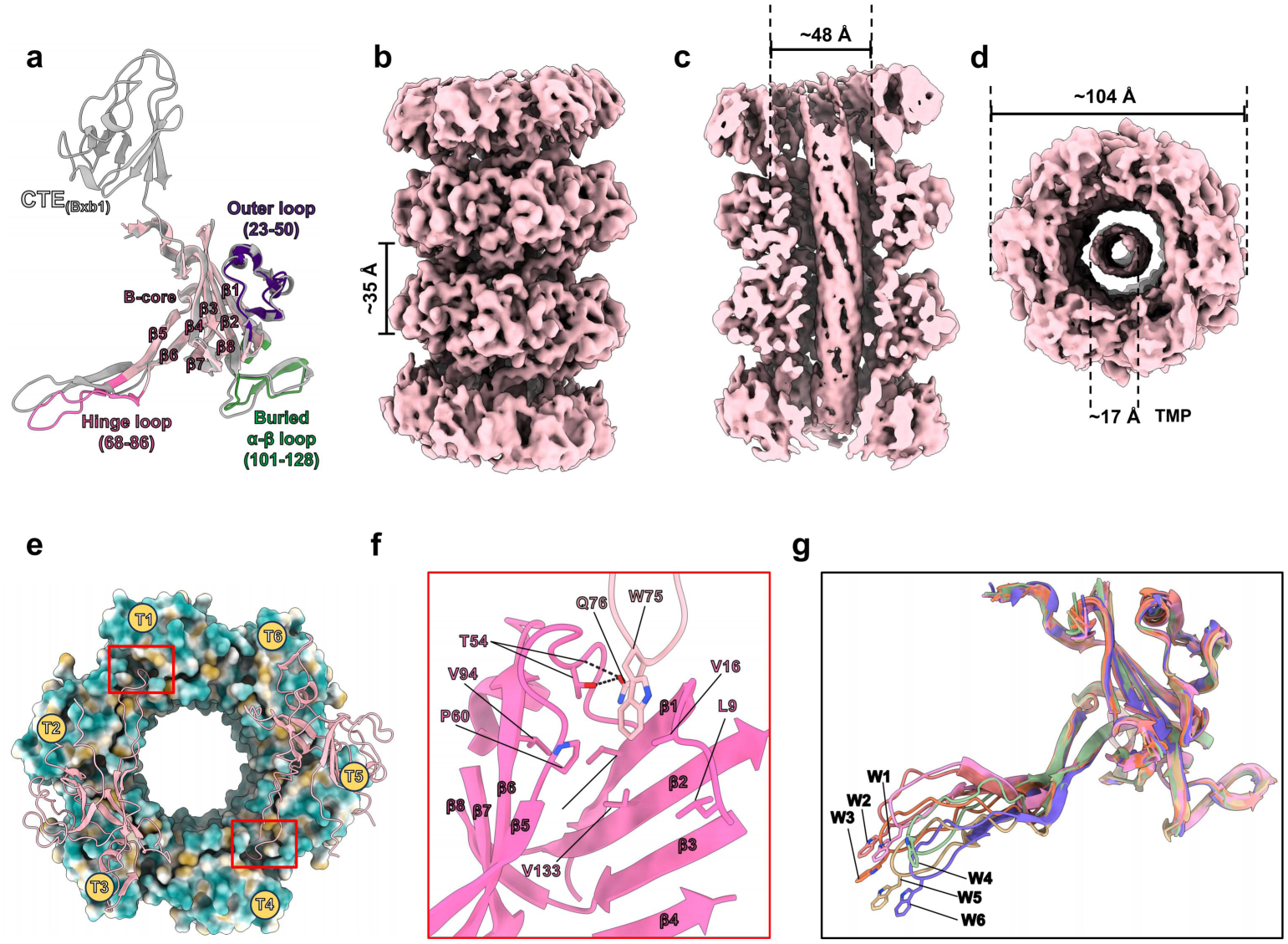
Structure and assembly of the D29 tail tube. **a**, Cartoon representation of a gp23 subunit, showing the β-core, outer loop, hinge loop and buried α–β loop. The C-terminal domain of Bxb1 gp12 is shown in gray for comparison. **b**, Cryo-EM map of four stacked gp23 rings, with an axial rise of approximately 35 Å between adjacent rings. **c**, Cutaway view of the tail tube, showing the approximately 48Å-wide central channel. **d**, Axial view of the C6 gp23 ring, showing its approximately 104Å outer diameter and the central density assigned to TMP. **e**, Hydrophobic surface representation of a gp23 hexamer, with the six subunits designated T1–T6. **f**, Cartoon representation of the boxed region in **e**, showing the insertion of the hinge loop into the hydrophobic pocket of an adjacent gp23 subunit. Hydrogen bonds are indicated by dashed lines. **g**, Structural superposition of the six gp23 subunits, highlighting conformational variation in their hinge loops and the positions of W75.

**Supplementary Fig. 9.**
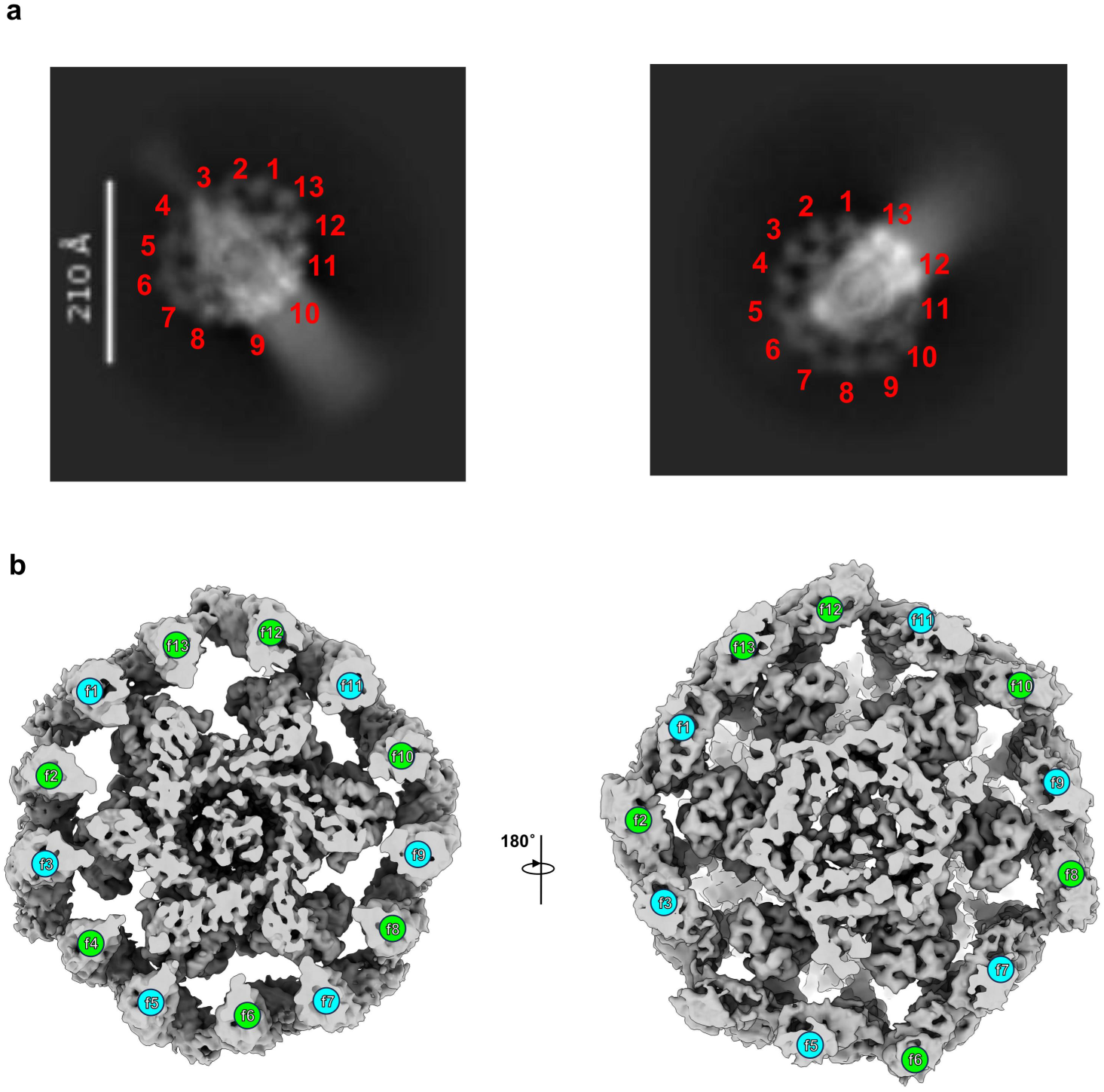
13-copy organization of the D29 gp6 collar. **a**, Representative 2D class averages of the D29 baseplate viewed along the tail axis, showing 13 peripheral densities corresponding to gp6. The densities are numbered 1-13. **b**, Axial views of representative C1 cryo-EM maps showing the 13 gp6 subunits (f1-f13) surrounding the central baseplate core.

**Supplementary Fig. 10.**
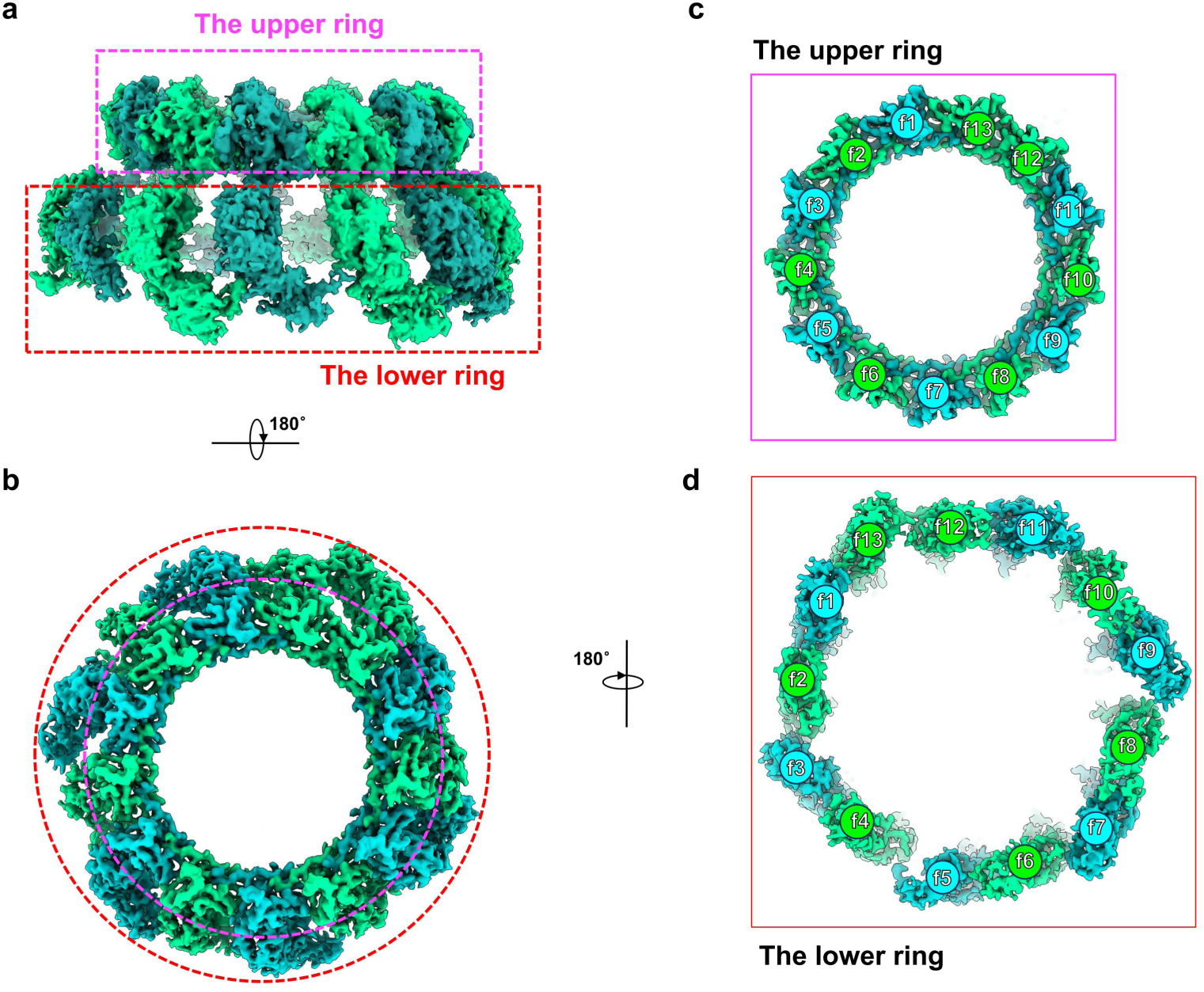
Organization of the gp6 baseplate layers. **a**, Side view of the 13-subunit gp6 collar assembly, with its upper and lower regions outlined by magenta and red dashed boxes, respectively. **b**, Axial view of the gp6 collar assembly viewed from the lower-region side, with the upper and lower regions indicated by magenta and red dashed circles, respectively. **c-d**, Isolated axial views of the pseudo-C13-symmetric upper ring (**c**) and asymmetric lower region (**d**) of the gp6 collar.

**Supplementary Fig. 11.**
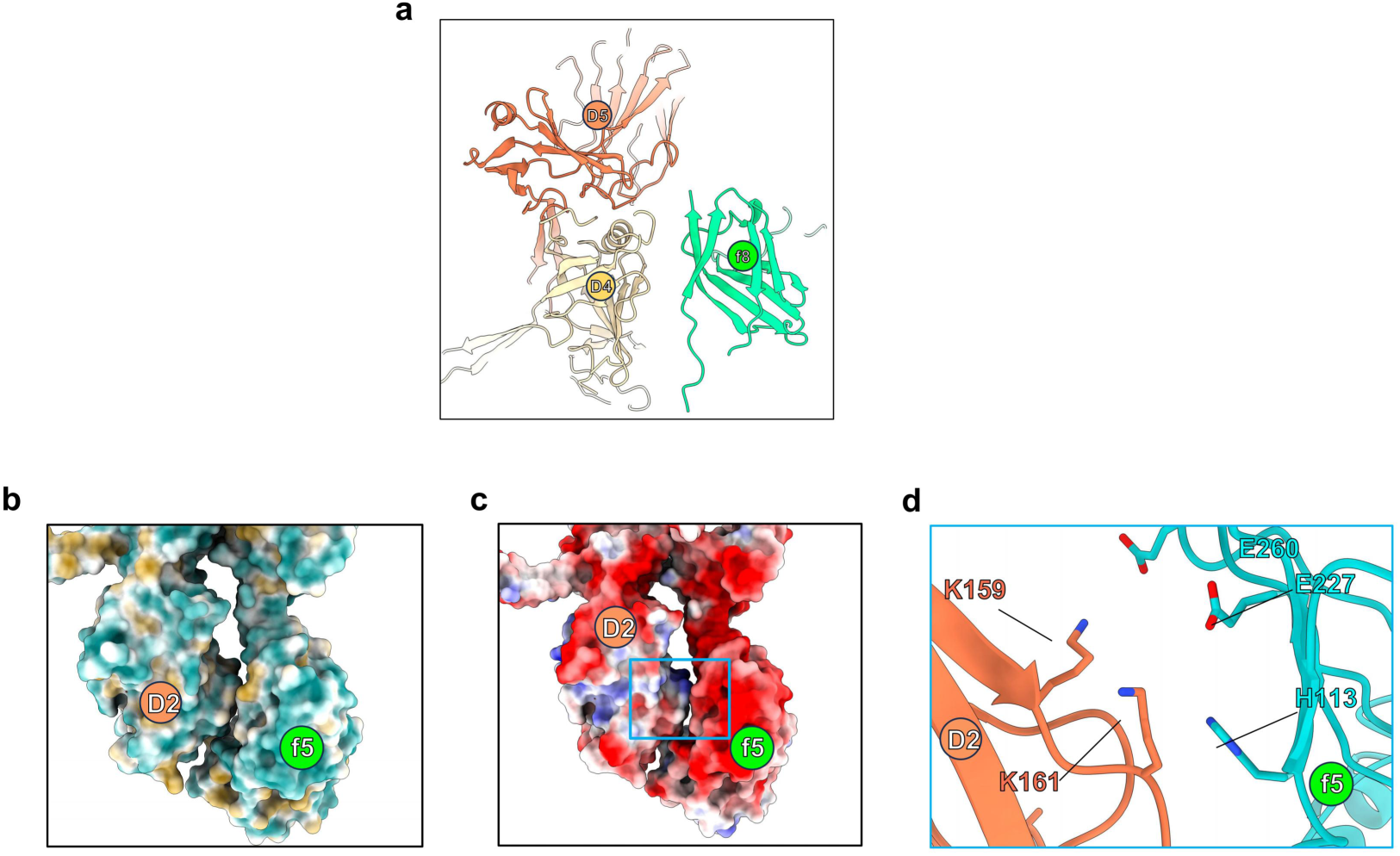
Limited contacts between gp27 and peripheral gp6 subunits. **a**, Cartoon representation showing gp6 subunit f6 positioned between gp27 subunits D4 and D5, without apparent direct contacts. **b-c**, Surface representations of the lower-region interface between gp27 subunit D2 and gp6 subunit f5, colored by hydrophobicity (**b**) and electrostatic potential (**c**). Red and blue indicate negative and positive electrostatic potentials, respectively. **d**, Close-up of the boxed region in c, showing selected charged and polar residues at the interface.

**Supplementary Fig. 12.**
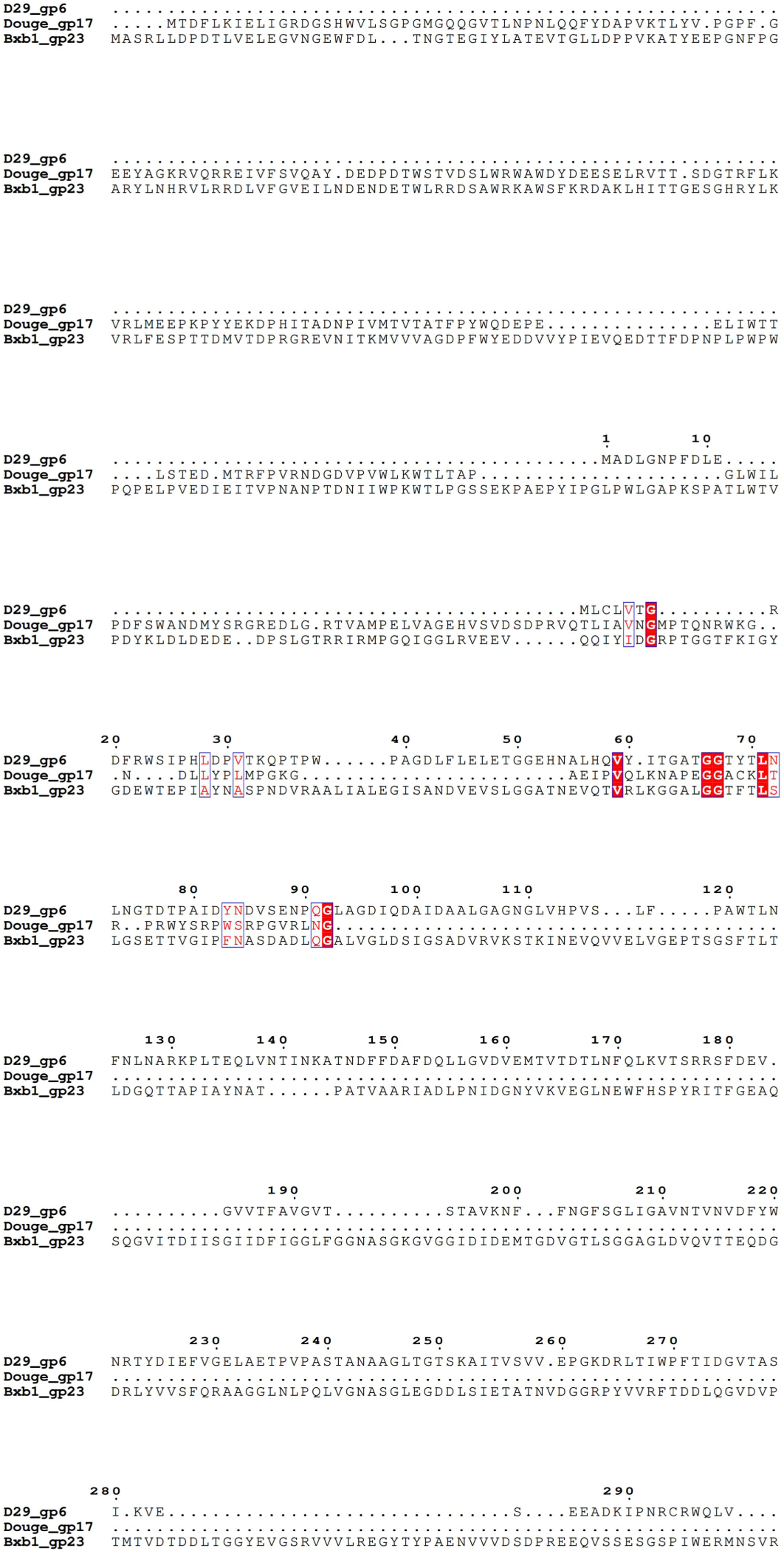
Multiple sequence alignment of D29 gp6, Douge gp17, and Bxb1 gp23.

**Supplementary Fig. 13.**
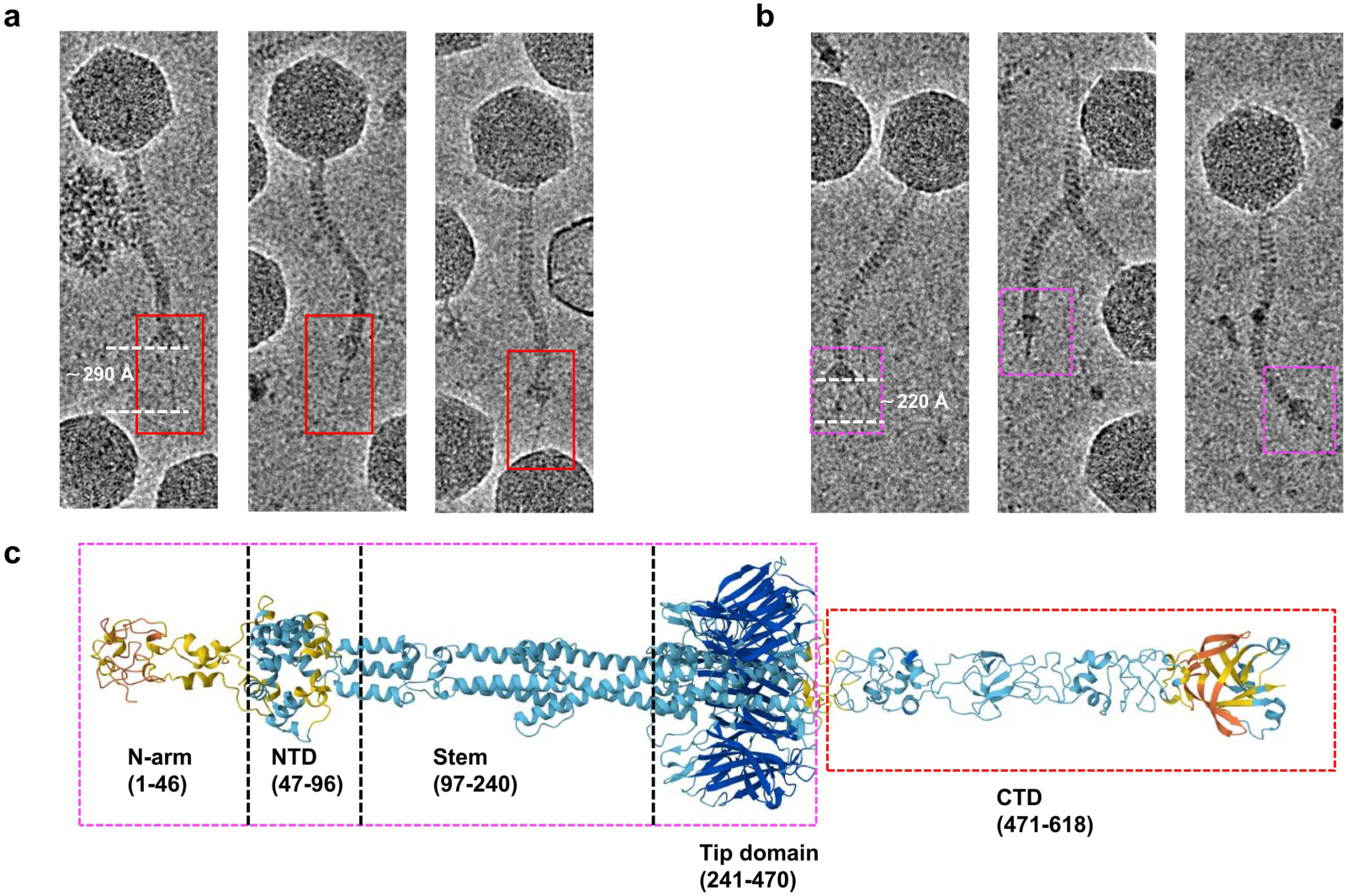
Structural variability and predicted domain organization of the D29 central fiber gp31. **a**, Representative cryo-EM micrographs of D29 particles showing an extended gp31 with an apparent length of approximately 290 Å. **b**, Representative cryo-EM micrographs showing the more frequently observed gp31 central fiber with an apparent length of approximately 220 Å. The gp31 fibers are highlighted by dashed boxes. **c**, AlphaFold3-predicted structure of the full-length gp31 trimer.

